## Supplemental Methods for "Crowdsourced identification of multi-target kinase inhibitors for RET- and TAU-based disease: the Multi-Targeting Drug DREAM Challenge"

**Supplemental File: All Submissions and Methods Descriptions/Writeups**

**Submission 9662708**

**Author: David Koes**

**Affiliation: University of Pittsburgh**

Prediction Methods:

Using convolutional neural network to score pharmacophore aligned poses with limited pose sensitivity. Representative structures for the desired targets were manually selected from the PDB and aligned. High resolution, ligand-bound, human protein structures were preferred. At most 3 structures were selected for each target.

For Problem 1

ret	2ivu 2x2k 5fm2 5amn

braf 5ita 5vam

src	1y57 2src

s6k	3wf8 3a60

mknk1 5wvd

ttk	4o6l 5n93

erk8 no structures

pdk1 5lvl 3nax

pak3 6fd3

For Problem 2

aurora 4uyn 4j8m 3fdn

pak1	4oth 5dew

fgfr1 4uwc 4uxq

lkb1	no bound structures

pak3	6fd3

tak1	5v5n 2yiy

pik3ca 5dxt 5ubr

The ligand bound structures for the primary targets (i.e. ret and aurora/pak1) were used as the bases for a pharmacophore search. The receptor and ligand were uploaded to the Pharmit search engine (http://pharmit.csb.pitt.edu), which automatically identified an interaction pharmacophore. The radius of all features was set to 1A. If the default pharmacophore was too specific, features were manually removed until the query returned a reasonable number of hits (at least in the hundreds). Solvent exposed features were preferentially removed. If the query was two broad, direction constrains were selectively added to hydrogen bond features to narrow the scope of the query. The MolPort library of 7 million commericially available compounds was searched. All the hits matching the query in addition to structures minimized in pharmit (using the AutoDock Vina scoring function) were downloaded.

These poses were then scored against all the protein structures using a gnina (http://github.com/gnina) and a 3D convolutional neural network that was trained on the PDBbind refined set to predict binding affinities. The model used here was trained only on reasonable docked poses and so has not learned to penalize steric clashes.

Scored compounds were then filtered to create a ranked list of compounds deemed responsive to the challenge. Affinity thresholds were manually tuned to select compounds with the desired relationships. Chemical novelty is assessed by comparing to the ChEMBL activity sets corresponding to the protein targets. The maximum Tanimoto (as calculatd by the default RDKit fingerprints) to _any_ ChEMBL compound in these sets is recorded. Compounds with Tanimoto > 0.7 are excluded.

Note that ten solutions are provided, in ranked order, in case there are difficulties acquiring the first five or there is sufficient budget to acquire additional compounds.

Rationale:

Why is your approach innovative?: The approach uses a convolutional neural network trained on a direct representation of the 3D structure of a protein-ligand complex to predict binding affinities. Such 3D deep learning approaches to structure based drug design are relatively new, having only emerged in the last few years, and have yet to be subjected to extensive prospective evaluation, as enabled by the DREAM challenge. This particular submission is unique in that we are explicitly using a model that is tolerant of steric clashes and (hopefully) will correctly score pharmacophore aligned poses without refining the ligand position.

Why will your approach be generalizable?: This approach builds upon established structure-based drug design workflows and can be applied to any target with a known receptor.

Problem 1:

- Solution 1:

ZINC ID: ZINC000067911340

VENDOR ID: MolPort-005-945-291

SMILES string: COc1cc(c2c(c1)oc(c(c2=O)c1c(cc(c2c1oc(cc2=O)c1ccc(cc1)O)O)O)c1ccc(cc1)O)O

VENDOR NAME: MolPort

Explanation of chemical novelty: Tanimoto=0.596477 src=7.202710 pak3-1=8.865550 pdk1-1=7.980150 s6k=7.443580 mknk1-3=8.007800 ttk-1=8.253620 ret=9.071110 braf=7.707130

- Solution 2:

ZINC ID: ZINC000096085369

VENDOR ID: MolPort-039-338-970

SMILES string: COc1ccc(cc1)[C@@H]1[C@H]([C@@H]2C(=O)c3c(cc(cc3O[C@@H]2c2ccc(cc2)O)O)O)C(=O)c2c(cc(cc2O1)OC)O

VENDOR NAME: MolPort

Explanation of chemical novelty: Tanimoto=0.545068 src=7.819430 pak3-1=8.638160 pdk1-1=8.557120 s6k=7.661320 mknk1-3=7.459990 ttk-1=8.784440 ret=8.861160 braf=7.589300

- Solution 3:

ZINC ID: ZINC000095913548

VENDOR ID: MolPort-039-338-895

SMILES string: COc1cc(c2c(c1)O[C@@H]([C@@H](C2=O)[C@H]1[C@@H](c2ccc(cc2)O)Oc2cc(cc(c2C1=O)O)O)c1ccc(cc1)O)O

VENDOR NAME: MolPort

Explanation of chemical novelty: Tanimoto=0.545997 src=7.677360 pak3-1=8.437050 pdk1-1=8.095580 s6k=7.439030 mknk1-3=7.260620 ttk-1=8.574550 ret=8.624020 braf=7.485800

- Solution 4:

ZINC ID:

VENDOR ID: MolPort-042-645-765

SMILES string: CC[C@H](C)[C@H](C(=O)Nc1cc2c(cc1)[nH]c(CCc1ccccc1)n2)Nc1ccc2-c3c(CC[C@@H](c2cc1=O)NC(=O)C)cc(c(c3OC)OC)OC

VENDOR NAME: MolPort

Explanation of chemical novelty: Tanimoto=0.674571 src=6.941540 pak3-1=7.525330 pdk1-1=7.669240 s6k=7.489600 mknk1-3=7.387870 ttk-1=8.401290 ret=8.427920 braf=6.486890

- Solution 5:

ZINC ID:

VENDOR ID: MolPort-019-936-923

SMILES string: CC(=CCC[C@@](C)([C@@H]1CC[C@@]2([C@H]1[C@@H](C[C@H]1[C@]2(C[C@@H]([C@@H]2[C@]1(CC[C@H](C2(C)C)O)C)O[C@H]1[C@@H]([C@H]([C@@H]([C@@H](CO)O1)O)O)O[C@H]1[C@@H]([C@H]([C@@H](CO1)O)O)O)C)O)C)O)C

VENDOR NAME: MolPort

Explanation of chemical novelty: Tanimoto=0.419698 src=8.352460 pak3-1=8.185400 pdk1-1=7.660210 s6k=7.981660 mknk1-3=7.116460 ttk-1=8.042210 ret=8.351780 braf=8.339820

- Solution 6:

ZINC ID:

VENDOR ID: MolPort-044-521-040

SMILES string: CCNS(=O)(=O)c1ccccc1S(=O)(=O)Nc1ccc2c(c1)CN(C)C(=O)N2

VENDOR NAME: MolPort

Explanation of chemical novelty: Tanimoto=0.509615 src=7.215470 pak3-1=7.954990 pdk1-1=8.069670 s6k=7.133740 mknk1-3=7.274000 ttk-1=6.911060 ret=8.286370 braf=7.603270

- Solution 7:

ZINC ID: ZINC000000354329

VENDOR ID: MolPort-000-279-564

SMILES string: c1(cc(cc(c1O)C(C)C)NS(=O)(=O)c1c(N)cccc1)C(C)C

VENDOR NAME: MolPort

Explanation of chemical novelty: Tanimoto=0.435530 src=7.044640 pak3-1=7.428440 pdk1-1=7.496350 s6k=6.663220 mknk1-3=6.509820 ttk-1=7.015780 ret=8.122230 braf=6.804350

- Solution 8:

ZINC ID: ZINC000009377085

VENDOR ID: MolPort-005-795-453

SMILES string: Nc1n[nH]c(CCCCCNC(=O)c2ccc(cc2)S(=O)(=O)Nc2ccc(cc2)F)c1C#N

VENDOR NAME: MolPort

Explanation of chemical novelty: Tanimoto=0.530355 src=6.844180 pak3-1=7.276990 pdk1-1=7.018430 s6k=6.434700 mknk1-3=6.888610 ttk-1=7.281150 ret=7.976400 braf=6.376420

- Solution 9:

ZINC ID: ZINC000096115072

VENDOR ID: MolPort-028-855-651

SMILES string: c1(c2cnn(c2nc2c1oc1c(C)c(O)ccc21)C(C)C)c1cc(c(c(c1)O)O)OC

VENDOR NAME: MolPort

Explanation of chemical novelty: Tanimoto=0.663972 src=6.690050 pak3-1=7.308890 pdk1-1=7.063310 s6k=6.097180 mknk1-3=6.745940 ttk-1=6.839280 ret=7.889260 braf=6.707610

- Solution 10:

ZINC ID:

VENDOR ID: MolPort-045-915-586

SMILES string: COc1cc2ccc1OCC(=O)NCCCNCCCCNC(=O)[C@@H]([C@H](C)O)NC(=O)[C@@H](CC(C)C)NC(=O)CC2

VENDOR NAME: MolPort

Explanation of chemical novelty: Tanimoto=0.618286 src=6.519400 pak3-1=6.886070 pdk1-1=7.048960 s6k=6.026080 mknk1-3=5.821110 ttk-1=7.132460 ret=7.594810 braf=5.971040

Problem 2:

- Solution 1:

ZINC ID: ZINC000150340658

VENDOR ID: MolPort-016-580-907

SMILES string: Cc1cc(=O)oc2c1ccc(c2)NC(=O)[C@H](CCC(=O)O)NC(=O)[C@H](CC(C)C)NC(=O)[C@H](CC(C)C)NC(=O)OCc1ccccc1

VENDOR NAME: MolPort

Explanation of chemical novelty: Tanimoto=0.560727 pak3-3=7.413280 aurora=8.564800 tak1-3=7.968470 pik3ca-1=6.844350 pak1=9.232770 fgfr1=7.899540

- Solution 2:

ZINC ID: ZINC000032927046

VENDOR ID: MolPort-009-641-699

SMILES string: CC(C)N(CCNS(=O)(=O)c1ccc(c(c1)[N+](=O)[O-])NCc1cc[nH]n1)C(C)C

VENDOR NAME: MolPort

Explanation of chemical novelty: Tanimoto=0.540444 pak3-3=5.711590 aurora=6.367070 tak1-3=5.787760 pik3ca-1=5.377990 pak1=6.558820 fgfr1=6.141540

- Solution 3:

ZINC ID:

VENDOR ID: MolPort-045-915-960

SMILES string: COCCCN1Cc2cc(c(cc2)OC)Oc2cc(CO[C@@H]3CCN(C[C@@H]3NC(=O)C1)C(=O)c1ccc3c(c1)nn[nH]3)ccc2

VENDOR NAME: MolPort

Explanation of chemical novelty: Tanimoto=0.528239 pak3-3=5.318060 aurora=6.219700 tak1-3=5.418300 pik3ca-1=5.942040 pak1=6.834830 fgfr1=5.960940

- Solution 4:

ZINC ID: ZINC000009509579

VENDOR ID: MolPort-004-094-563

SMILES string: COc1ccccc1NC(=O)c1ccc(cc1)NC(=O)CCN1C(=O)c2ccccc2C1=O

VENDOR NAME: MolPort

Explanation of chemical novelty: Tanimoto=0.505387 pak3-3=5.512520 aurora=6.161180 tak1-3=5.326870 pik3ca-1=4.777780 pak1=6.596950 fgfr1=5.724830

- Solution 5:

ZINC ID: ZINC000029828001

VENDOR ID: MolPort-009-593-551

SMILES string: COc1ccc(CCN(C)C(=O)CSc2nnc(Nc3cccc(C)c3)s2)cc1OC

VENDOR NAME: MolPort

Explanation of chemical novelty: Tanimoto=0.408115 pak3-3=5.049610 aurora=5.811150 tak1-3=5.086820 pik3ca-1=4.372320 pak1=5.624710 fgfr1=5.752320

- Solution 6:

ZINC ID:

VENDOR ID: MolPort-028-730-964

SMILES string: O=C(c1cccc(c1)S(=O)(=O)NCc1cc[nH]n1)NC[C@H]1OCCOC1

VENDOR NAME: MolPort

Explanation of chemical novelty: Tanimoto=0.427032 pak3-3=4.898510 aurora=5.751490 tak1-3=5.076800 pik3ca-1=4.691640 pak1=5.719520 fgfr1=5.946270

- Solution 7:

ZINC ID: ZINC000009884489

VENDOR ID: MolPort-005-328-937

SMILES string: CCOC(=O)c1c(NC(=O)CSc2nnc(C)c(=O)n2N)sc(C)c1C

VENDOR NAME: MolPort

Explanation of chemical novelty: Tanimoto=0.553655 pak3-3=5.119570 aurora=5.666640 tak1-3=5.052490 pik3ca-1=4.229370 pak1=6.479960 fgfr1=5.172320

- Solution 8:

ZINC ID: ZINC000096309558

VENDOR ID: MolPort-044-560-719

SMILES string: C[C@@H]1[C@H]([C@@H]([C@H]([C@H](O1)O[C@@H]1[C@@H](CO)O[C@@H]([C@@H]([C@@H]1O)O)O[C@H]([C@@H](CO)O)[C@@H]([C@H](C=O)O)O)O)O)N[C@H]1C=C([C@H]([C@@H]([C@H]1O)O)O)CO

VENDOR NAME: MolPort

Explanation of chemical novelty: Tanimoto=0.486757 pak3-3=3.720480 aurora=5.122080 tak1-3=4.598510 pik3ca-1=3.909720 pak1=5.814780 fgfr1=4.588220

- Solution 9:

ZINC ID:

VENDOR ID: MolPort-001-741-535

SMILES string: CC(=O)N[C@@H]1[C@H]([C@@H]([C@@H](CO[C@H]2[C@@H]([C@H]([C@@H](CO2)O[C@@H]2[C@@H]([C@](CO2)(CO)O)O)O[C@@H]2[C@H]([C@@H]([C@@H](CO2)O)O)O)O)O[C@H]1O[C@H]1CC[C@]2([C@H](C1(C)C)CC[C@@]1([C@H]2CC=C2[C@]1(C[C@@H]([C@@]1([C@H]2CC(CC1)(C)C)C(=O)O[C@@H]1[C@H]([C@@H]([C@@H](CO1)O)O)O)O)C)C)C)O)O

VENDOR NAME: MolPort

Explanation of chemical novelty: Tanimoto=0.560634 pak3-3=4.130950 aurora=5.012430 tak1-3=3.764290 pik3ca-1=4.017180 pak1=6.138050 fgfr1=4.267020

- Solution 10:

ZINC ID: ZINC000075352807

VENDOR ID: MolPort-010-846-490

SMILES string: CCOC(=O)c1c(C)[nH]c(CCC(=O)NCCCN2CCOCC2)c1C

VENDOR NAME: MolPort

Explanation of chemical novelty: Tanimoto=0.484448 pak3-3=4.261170 aurora=4.908140 tak1-3=4.301910 pik3ca-1=4.085280 pak1=5.825020 fgfr1=4.984630

**Submission 9662305**

**Author: Team Stratified (4 anonymous members)**

Prediction Methods: |

Due to technical advances more and more profiling data becomes available, especially in the field of kinase research [1] and machine learning algorithms can be applied to a broader range of drug discovery challenges. However, most of the available data is highly imbalanced and biased towards inactive compounds. A major issue is that machine learning on skewed class labels tends to result in models with low sensitivity due to overfitting.

We aimed to develop a model that, firstly, addresses this problem by using gradient boosting models of the LightGBM library [2] to increase the loss of false negative predictions. Cross-validation model performance can be deceiving when random sampling from highly correlated data sets [4]. A model might perform well either due to good generalization or by pure chance because of highly correlated data between folds. To prevent this, we developed a novel stratified cluster sampling strategy that leads to low correlation between folds while simultaneously preserving the underlying class (active/inactive) distribution (manuscript in preparation). This sampling method combined with Bayesian Optimization [5,6] was used to find an optimal set of hyperparameters for each target endpoint. The mean weighted F1 score based on a nested 5-fold stratified cluster cross-validation was introduced as the parameter evaluation metric.

Secondly, since some targets are still underexplored and have relatively little assay data available (especially too few active compounds), we decided to build multi-target models for the groups of desired on- and off-targets (see details below). Such strategies have been shown to be beneficial, especially when the targets of interest are related which holds true for kinases.[3]

A combination of publicly available data sets [7] was used to train and evaluate the gradient boosting models. A pIC50 value of 6.3 was used as activity threshold, compounds with activities above this value are considered active on a specific target. The compounds were standardized and represented by a vector containing the ECFP4 fingerprints of size 4096 and 111 molecular descriptors both calculated using the RDKit library. For the two challenges, the following data was collected (in parenthesis number of active compounds/ number of inactive compounds using the above-mentioned threshold).

Challenge 1 (Ch1)

- Required on-target RET (387/1187)

- Required off-target MKNK1 (46/623)

- Desired on-targets BRAF (1177/854), SRC (1019/1928), S6K (620/1253)

- Desired off-targets TTK (299/764), Erk8 (68/216), PDK1 (186/1535), PAK3 (25/640)

- Merged groups BRAF_group (BRAF, SRC, S6K); MKNK1_group (MKNK1, TTK, Erk8, PDK1)

Challenge 2 (Ch2)

- Required on-target PAK1 (103/1385), AURKA (1449/2024)

- Required off-target PAK3 (25/640), TAK1 (54/289)

- Desired on-targets FGFR1 (448/1819), LKB1 (16/294)

- Desired off-targets PIK3CA (no data available)

- Merged groups PAK3_group (PAK3, TAX1); LKB1_group (FGFR1, LKB1)

For each endpoint (which can be a single target or a target group) an ensemble model is created based on the resulting five hyperparameter sets found by the optimization routine. The outer-fold performance from the cross-validation of the resulting ensembles showed promising AUC values, i.e., RET 0.83 (+-0.03), MKNK1_group 0.75 (+-0.07), BRAF_group 0.94 (+-0.00), PAK1 0.94 (+-0.04), AURKA 0.89 (+-0.00), PAK3_group 0.79 (+-0.17) and FGFR1_group 0.90 (+-0.04).

With the described machine learning models in hand, a total of 13 million compounds from the ZINC15 database were screened. Potential leads were ranked by a scoring formula combining the resulting posterior probabilities of the individual models. The probabilities range between 0 and 1, the higher the value the higher the probability that the compound binds to the respective target. In the combined score, the probabilities of the required targets are up-weighted by a factor of two and the desired target probabilities are used as is (factor of 1). The probabilities for on-targets are summed-up, while the values for off-targets are subtracted from the total score.

After the ML-based screening and ranking, the top 1000 compounds were further filtered based on novelty and drug-likeness. First, compounds are discarded if their tanimoto similarity to known inhibitors for the respective required on- and off-target equals one. Second, compounds are dropped if they violate more than one rule of Lipinski’s rule of five.

The final set of high ranking compounds were finally screened against the required targets using structure-based methods (licensed software) and manually checked for their fit in the binding site of the respective required target.

Rationale:

Why is your approach innovative?: |

Nowadays that more and more profiling data becomes available, machine learning algorithms provide a valuable tool to solve drug discovery challenges. We developed a novel stratified clustering algorithm to account for correlation in the compound data and to achieve well regularized models using bayesian hyperparameter optimization. Since still not every kinase has enough data, be built multi-target models, exploiting the fact that kinase binding sites are relatively conserved. Furthermore, with our combined scoring scheme, we prioritize compounds that have a high chance to bind to the on-targets, while penalizing high binding probabilities to known off-targets. Thus, our method takes several targets into account, while allowing for fast screening of large data sets such as ZINC15. The calculated probability scores do neither dependent on available protein structures nor on information about different conformations such as DFG-in or DFG-out.

For the final selection from the top 1000 compounds, nevertheless, we included information from structure-based methods to validate the actual fit of the compounds in the binding site of the required on-target.

Why will your approach be generalizable?: |

Our machine learning method can be applied to any multi-target drug design project for which a decent amount of profiling data is available (more than 50 data points for each class). Furthermore, since kinases are highly similar, combining single target models to multi-target models for sets of on- and off-targets can help to improve the performance. Our ranking equation can be adapted to individual preferences for specific on- or off-targets based on the weighting factor, in the scoring function on-targets probabilities are accounted as positive contributions, off-targets probabilities as negative values. This procedure can be applied to any other combination of multi-kinase drugs.

Problem 1:

- Solution 1:

ZINC ID: ZINC000057510750

VENDOR ID: MolPort-010-816-078

SMILES string: Cc1ccc(Nc2n[nH]nc2C(=O)Nc2ccc3c(c2)OCCO3)cc1C

VENDOR NAME: MolPort

Explanation of chemical novelty: Similar ZINC compounds with good scores ZINC000057510743. Low tanimoto similarity to known RET (0.33) or other kinase inhibitors. Probability to bind to RET 0.71, MKNK1_group 0.18.

- Solution 2:

ZINC ID: ZINC000205928292

VENDOR ID: MolPort-044-830-664

SMILES string: CC(C)n1nc(-c2ccc(NC(=O)Nc3cc(C(F)(F)F)ccc3F)cc2)c2c(N)ncnc21

VENDOR NAME: MolPort

Explanation of chemical novelty: Similar ZINC compounds with good scores, e.g., ZINC000002576348, ZINC000013132888, ZINC000036056301. Low tanimoto similarity to known RET inhibitors (0.42). Probability to bind to RET 0.83, MKNK1_group 0.21.

- Solution 3:

ZINC ID: ZINC000072117895

VENDOR ID: MolPort-020-184-322

SMILES string: Cc1cc(N2CCCC2)nc(Nc2ccc(NC(=O)Nc3ccc(C)c(C)c3)cc2)n1

VENDOR NAME: MolPort

Explanation of chemical novelty: Cluster of this compound serious with good scores, e.g., ZINC000064801478, ZINC000072437051, ZINC000064801495, ZINC000072117563, ZINC000021795483, all in stock at MolPort, not yet used in kinase context. Low tanimoto similarity to known RET (0.29) or other kinase inhibitors. Probability to bind to RET 0.74, MKNK1_group 0.20.

- Solution 4:

ZINC ID: ZINC00004176475

VENDOR ID: C200-0356

SMILES string: Cc1ccc(NC(=O)c2c(C)nc3sc(C(N)=O)c(N)c3c2-c2ccco2)c(C)c1

VENDOR NAME: ChemDiv

Explanation of chemical novelty: Preferred compound is ZINC00004176475. If not in stock use analogues such as ZINC000008592148 (MolPort-007-595-272) or ZINC000008592149 which are slightly too large to fulfill ¾ Lipinski's rule of 5. Low tanimoto similarity to known RET (0.31) or other kinase inhibitors. Probability to bind to RET 0.76, MKNK1_group 0.20.

- Solution 5:

ZINC ID: ZINC000033009328

VENDOR ID: MolPort-007-808-595

SMILES string: Cc1ccc(C(=O)Nc2ccc(N3CCCC3)c(NC(=O)Nc3ccc(C)cc3Cl)c2)cc1

VENDOR NAME: MolPort

Explanation of chemical novelty: Low tanimoto similarity to known RET (0.30) or other kinase inhibitors. Probability to bind to RET 0.73, MKNK1_group 0.21.

Problem 2:

- Solution 1:

ZINC ID: ZINC000257236672

VENDOR ID: LAS34152690

SMILES string: Cc1cc2nc(CCC(=O)N(C)C[C@@]3(O)CCCN(c4ccnc5cccnc45)C3)[nH]c2cc1C

VENDOR NAME: Asinex

Explanation of chemical novelty: Low tanimoto similarity to known Pak1 and AurA inhibitors. Probability to bind to Pak1 0.58, AurA 0.55, Pak3_group 0.45. Note that similar (as well as different) compounds with higher Pak1 and AurA probability were also present, such as ZINC000257272595 (AurA 0.69, Pak1 0.62, Pak3 0.52), but need further time for evaluation. Generally, Pak1 and Pak3 share high sequence similarity and it is difficult to distinguish between these two kinases. Please note that due to time reasons we could not fully finish the selection for challenge 2 compounds and would appreciate if you could put more weight on challenge 1 results.

**Submission 9662301**

**Authors: AC Tan^1^, Jaewoo Kang^2^, Minji Jeon^2^, Jinhyuk Lee, Hwisang Jeon, Miyoung Ko, Donghyeon Park^2^**

**Affiliations: ^1^University of Colorado Anschutz Medical Campus,^2^ Korea University**

Prediction Methods: We made a prediction of ZINC compounds through a total of three steps. The description of the three steps are as follows.<Step 1 Kinase inhibitor candidate extraction> KIEO(tanlab.ucdenver.edu/KIEO/KIEOv1.0/) is KInase Experiments Omnibus, developed by one of our team members, Professor Aik-Choon Tan of the University of Colorado. The KIEO database was constructed by collecting and curating published experimental data of kinase inhibitors from more than 600 articles. We utilized KIEO to identify kinase inhibitor candidates that satisfy the conditions given in Subchallenge 1 and 2. Some of our answers include the kinase inhibitors found in Step 1 (Figure 1A is available at infos.korea.ac.kr/dmis_mtd) <Step 2 Choosing a ZINC chemical compound candidates with a response similar to the kinase inhibitor candidates via the Cmap score predictor? We designed a deep-learning based Cmap score predictor. The Cmap score is a similarity score between two compounds obtained from the perturbation-driven gene expression dataset, called Connectivity map(Cmap), provided by Broad institute[1]. We can extract the Cmap scores of 2.8M compound pairs of 2400 single compounds. Top 5% and bottom 5% of the cmap score are used for our deep learning model. Our model is a classification model that is predicting True and False labels representing similarities for each pair of compounds. The input of the model is pairs of fingerprints of two chemical compounds. Once input pairs and corresponding scores (Cmap scores) are given, we feed each input pair to our Siamese network. If trained properly, the Siamese network is able to predict Cmap scores of unseen input pairs where one of the fingerprints (or both of them) is not seen during training. Overview of our model is illustrated in Figure 1B which is available at infos.korea.ac.kr/dmis_mtd. Motivated by the work of Koch [2], we share the weights of two multilayer perceptrons to build a Siamese network. We feed two different inputs to two separate MLPs that share weights, and gather the outputs of each MLP. Outputs of the MLPs are used for predicting the Cmap scores of the two fingerprint inputs. We compute the weighted L2 distance of two outputs, and using sigmoid function, the distance is transformed into a probability of classifying Cmap score as 1. For MLPs, we used 3 fully connected layers with hidden dimension set to 100 and sigmoid nonlinearity between the layers. For evaluating our model, we set up a real world experimental environment where responses of some drugs are known, but some are not. We wanted to know the similarity between well-known and unknown drugs, or between unknown drugs. Therefore we randomly divided the drugs in Cmap into 90% and 10%. 10% of the drugs were excluded from the training set and included only in the validation set and test set. Our Siamese Network achieved 0.755 of F1 score in all test pairs with 0.866, 0.694, 0.647 of F1 scores for the pairs of the known drugs and the known drugs, the pairs of the known drugs and the unknown drugs, and the pairs of the unknown drugs and unknown drugs, respectively. For submitting to DREAM Challenge, we generated pairs of compounds to be tested. One pair is a combination of two compounds and each compound is from kinase inhibitor found in Step 1 and a chemical compound in ZINC chemical compounds. <Step 3> Filtering out ZINC chemical compound candidates through Lipinski's rules, patent search and protein-ligand docking prediction algorithm. Through the process up to step 2, we were able to select several ZINC chemical compound candidates. We checked Lipinski's rules, searched patents through PubChem, and considered predicted binding affinity using protein-ligand docking algorithm for filtering out ZINC chemical compound candidates (Figure 1C). ref 1. Subramanian, Aravind, et al. "A next generation connectivity map; L1000 platform and the first 1,000,000 profiles." Cell 171.6 (2017) ref 2. Koch, Gregory, Richard Zemel, and Ruslan Salakhutdinov. "Siamese neural networks for one-shot image recognition." ICML Deep Learning Workshop. Vol. 2. 2015.

Rationale:

Why is your approach innovative?: Our method is innovative in several ways. The first one is that we were able to find kinase inhibitors that meet all the criteria by searching on KIEO, KInase Experiment Omnibus. Second, we generated a deep learning-based Siamese network model. The model predicts the similarity of gene expression level effects using only the structural information of two compounds.

Why will your approach be generalizable?: Our Siamese network can be generalized because it performs well for drugs that have no knowledge of the reactivity. Siamese network is well known for its ability to predict the degree of similarities of unseen input pairs.

Problem 1:

- Solution 1:

ZINC ID: ZINC98209221

VENDOR ID: Sigma Aldrich(SML1332|SIGMA), Chem Scene(CS-0875), MedChem Express Economical(HY-15434), Bioactive Tocris(5429), Molport(MolPort-039-101-308, MolPort-042-665-858)

SMILES string: CCN1CCN(Cc2ccc(NC(=O)c3ccc(C)c(Oc4ccnc5[nH]ccc45)c3)cc2C(F)(F)F)CC1

VENDOR NAME: Sigma Aldrich, Chem Scene, MedChem Express Economical, Bioactive Tocris, Molport

Explanation of chemical novelty: ZINC98209221, also known as NG25, is originally developed as the TAK1 (MAP3K7) inhibitor. Through our KIEO database, it shows that NG25 has high binding activity against the targets of RET(M918T), BRAF, SRC and S6K (Percent of binding > 80%), and not binding to MKNK1, TTK, ERK8, PDK1 and PAK3. As the original intended target for this compound is TAK1, here, using our approach, we predict that this compound could be repurposed to inhibit the intended targets and avoiding the unintended targets.

- Solution 2:

ZINC ID: ZINC6745272

VENDOR ID: MedChem Express Economical(HY-10331, HY-10331A, HY-13308), Molport SC Economical(MolPort-009-679-472), Specs(AT-229/JK30570), Combi-Blocks(QC-8261), Matrix Scientific(120838, 147558)

SMILES string: CNC(=O)c1cc(Oc2ccc(NC(=O)Nc3ccc(Cl)c(C(F)(F)F)c3)c(F)c2)ccn1

VENDOR NAME: MedChem Express Economical, Molport SC Economical, Specs, Combi-Blocks, Matrix Scientific

Explanation of chemical novelty: ZINC6745272, also known as Regorafenib, is a FDA-approved small molecule multi-kinase inhibitor. From our prediction, this compound inhibits the targets with high specificity and not binds to the unintended targets.

- Solution 3:

ZINC ID: ZINC4916928

VENDOR ID: Molport SC Economical(MolPort-004-498-182), UORSY(PB17581104), eMolecules(11682036), Mcule(MCULE-3936899675)

SMILES string: C[C@H](OC(=O)CCNS(=O)(=O)c1ccc(F)c(Cl)c1)C(=O)NC(N)=O

VENDOR NAME: Molport SC Economical, UORSY, eMolecules, Mcule

Explanation of chemical novelty: According to our Cmap score predictor, ZINC4916928 is predicted to be similar to the kinase inhibitor candidates from KIEO. Moreover, we double-checked this compound through Lipinski's rules, patent search and protein-ligand docking prediction algorithm. We think this compound to be a new drug candidate for this subchallenge.

- Solution 4:

ZINC ID: ZINC4916940

VENDOR ID: UORSY(PB17581104), Molport SC Economical(MolPort-004-498-182), eMolecules(11682036), Mcule(MCULE-3936899675)

SMILES string: C[C@@H](OC(=O)CCNS(=O)(=O)c1ccc(F)c(Cl)c1)C(=O)NC(N)=O

VENDOR NAME: UORSY, Molport SC Economical, eMolecules, Mcule

Explanation of chemical novelty: According to our Cmap score predictor, ZINC4916940 is predicted to be similar to the kinase inhibitor candidates from KIEO. Moreover, we double-checked this compound through Lipinski's rules, patent search and protein-ligand docking prediction algorithm. We think this compound to be a new drug candidate for this subchallenge.

- Solution 5:

ZINC ID: ZINC328621612

VENDOR ID: Intermed(IMED1757490204), Enamine-REAL(Z1757408967), Molport(MolPort-039-176-182), eMolecules(112028455)

SMILES string: O=C(CCC1CCCCC1)N=c1ccn([C@H]2CCNC2)[nH]1

VENDOR NAME: Intermed, Enamine-REAL, Molport, eMolecules

Explanation of chemical novelty: According to our Cmap score predictor, ZINC328621612 is predicted to be similar to the kinase inhibitor candidates from KIEO. Moreover, we double-checked this compound through Lipinski's rules, patent search and protein-ligand docking prediction algorithm. We think this compound to be a new drug candidate for this subchallenge.

Problem 2:

- Solution 1:

ZINC ID: ZINC18279871

VENDOR ID: Molport BB Economical(MolPort-044-724-186), APExBIO(B6865), 1717 CheMall Corporation(HE005339HE285952HE315808), eMolecules(30487844), Tractus(TRA0029412)

SMILES string: O=C1N=c2ccccc2=C1c1[nH]c2ccccc2c1NO

VENDOR NAME: Molport BB Economical, APExBIO, 1717 CheMall Corporation, eMolecules, Tractus

Explanation of chemical novelty: ZINC18279871, also known as Indirubin-3'-monoxime, is originally developed as the CDK5 inhibitor. Through our KIEO database, it shows that NG25 has high binding activity against the targets of AURKA, PAK1, FGFR1, and LKB1, and low binding activity against PAK3, TAK1, and PIK3CA. As the original intended target for this compound is CDK5, here, using our approach, we predict that this compound could be repurposed to inhibit the intended targets and avoiding the unintended targets.

- Solution 2:

ZINC ID: ZINC103702671

VENDOR ID: 1717 CheMall Corporation(HE005462, HE345512)

SMILES string: CCCCc1c2c([nH]c1=C1C=CC(=O)C=C1)=NC=CN2

VENDOR NAME: 1717 CheMall Corporation

Explanation of chemical novelty: ZINC103702671 is also known as Aloisine A. Through our KIEO database, it shows that Aloisine A has high binding activity against the targets of AURKA, PAK1, FGFR1, and LKB1, and low binding activity against PAK3, TAK1, and PIK3CA.

- Solution 3:

ZINC ID: ZINC96998798

VENDOR ID: Molport SC Economical(MolPort-029-907-194), eMolecules(49291323), Mcule(MCULE-9099787337)

SMILES string: O=C(c1cccc([N+](=O)[O-])c1)c1nc2ccccc2[nH]1

VENDOR NAME: Molport SC Economical, eMolecules, Mcule

Explanation of chemical novelty: According to our Cmap score predictor, ZINC96998798 is predicted to be similar to the kinase inhibitor candidates from KIEO. Moreover, we double-checked this compound through Lipinski's rules, patent search and protein-ligand docking prediction algorithm. We think this compound to be a new drug candidate for this subchallenge.

- Solution 4:

ZINC ID: ZINC26776964

VENDOR ID: UORSY(PB354456136), Enamine-REAL(Z354378668), Molport SC Economical(MolPort-009-633-254), eMolecules(44087400), Mcule(MCULE-3249409325)

SMILES string: Fc1cccc(-c2nc(Cn3c(COc4ccccc4)nc4ccccc43)co2)c1

VENDOR NAME: UORSY, Enamine-REAL, Molport SC Economical, eMolecules, Mcule

Explanation of chemical novelty: According to our Cmap score predictor, ZINC26776964 is predicted to be similar to the kinase inhibitor candidates from KIEO. Moreover, we double-checked this compound through Lipinski's rules, patent search and protein-ligand docking prediction algorithm. We think this compound to be a new drug candidate for this subchallenge.

- Solution 5:

ZINC ID: ZINC12311718

VENDOR ID: UORSY(PB18421376), Molport SC Economical(MolPort-009-592-311), eMolecules(43721080), Mcule(MCULE-8421407902)

SMILES string: CCOC(=O)c1ccc(NC(=O)CSc2nc3cc([N+](=O)[O-])ccc3n2-c2ccccc2OC)cc1

VENDOR NAME: UORSY, Molport SC Economical, eMolecules, Mcule

Explanation of chemical novelty: According to our Cmap score predictor, ZINC12311718 is predicted to be similar to the kinase inhibitor candidates from KIEO. Moreover, we double-checked this compound through Lipinski's rules, patent search and protein-ligand docking prediction algorithm. We think this compound to be a new drug candidate for this subchallenge.

**Submission 9662288**

**Authors: Miguel Rocha^1^, Delora Baptista, Jorge Miguel Lourenço Ferreira^2^**

**Affiliations: ^1^Centre Biological Engineering, ^2^University of Minho**

Prediction Methods: A recent paper (doi 10.1021/acs.jcim.7b00316) describes a method (SEA+TC) which combines the Similarity Ensemble Approach with the maximum Tanimoto coefficient (maxTC) to predict binding affinity. In this paper, SEA+TC was performed for all of the compounds present in the ZINC15 database that are available for purchase. The resulting predictions for each target gene are available from the ZINC15 website, where only ligands which are highly similar (maxTC > 40) to ligands already known to bind to each of the targets are considered. \nThe approach we propose uses the SEA+TC predictions (p-value and maxTC) available in ZINC15 to select compounds that may have the ability to bind to multiple targets.\nWe used the Python language to develop a script to access the API from the ZINC15 database and to filter the results according to the criteria specified by the DREAM challenge.\nThe script accepts a list of targets and list of anti-targets as inputs, identified by their gene names. Lists specifiying required targets or required anti-targets can also be passed as inputs.\n\nFirst, we developed two different functions to obtain the information needed for our analysis. The first one accepts a list of genes and retrieves a matrix of the predictions for each gene which contains the four different columns, one has the ZINC ID (which identifies the compound), the second one containing the p-value, the third with the information of maxTc and the last one with the name of the gene being evaluated. This is obtained with the following request, "url = "http://zinc15.docking.org/genes/" + target + "/predictions/subsets/purchasable.json:gene_name+pvalue+maxtc+zinc_id?count=all"". This was used for both targets and anti-targers, for both challenge 1 and 2.\nFor further filtering of the results, we also built another function to retrieve information related to the compounds. With a similar approach to the previous funtion, we accessed the ZINC15 API in order to retrieve information about the compounds to filter the results even further. With the "url = "http://zinc15.docking.org/substances/" + zinc_id + ".json:mwt+hbd+hba+logp+tpsa"", we obtained the molecular weight (mwt), H-bond donors (hbd), H-bond acceptors (hba) and parametric Polar Surface Area (tSPA). The output for this function is a matrix where each row is a compound and the columns are the molecular features described.\n\nTherefore, for each of the targets and anti-targets, the respective SEA+TC predictions were retrieved from the ZINC15 database. The predicted ligands for the first required target was considered the starting point, and then the predicted ligands for any remaining required targets were evaluated against this initial set of ligands. Only the ligands that appeared in the SEA+TC predictions for all of the required targets were kept for the following steps. Afterwards, any ligands that belonged to the set of predictions for any of the required anti-targets were removed from the set of selected ligands. A simmilar approach was then applied to the desired targets, excluding compounds that have not been predicted to have binding affinity with the desired target. However, if none of the predicted ligands for a given desired target belonged to the set of previously selected ligands, the desired target was simply ignored. For desired anti-targets, any of the proposed ligands that also appear in the set of predictions for any of the desired anti-targets were excluded.\nTo further filter the set of selected ligands, we used the data available in ZINC15 for each compound to exclude ligands that did not adhere to Lipinski s rule of 5 and, for challenge 2, to guarantee that the proposed solutions had a polar surface area of less than 75.\nAs a final step, we decided to rank the obtained results using the mean of the p-value and maxTc. Since one of the requirements of this challenge is the requirement of RET (M918T) for challenge 1 and Aurora Kinase A, PAK1 for challenge two, we applied simple weight measures to meet this. So, we calculated the mean for all the p-values for the all the predicitions with a simple twist, the required genes have the weight of 1, where the other genes have the weight of 0.5. This is also applied to the maxTc values. With the means obtained, we ranked first the higher means for the p-value and lower one for the maxTc. We then calculated an average of the two ranks and selected the highest ones.

Rationale:

Why is your approach innovative?: Our approach is innovative in many ways. One of the main reasons is the fact that the approach used to make the new predictions for the ZINC database is relatively new (as stated before, the article is around one month old). Due to this, the huge amount of new data generated are still subject to new explorations. With the framework presented for this challenge, we take a naive yet strong approach to tackle the objectives purposed. Also, the framework explored, when taking into the account the SEA algorithm, it can be applied to other databases, enriching the process.

Why will your approach be generalizable?: Our approach is generalizable because the same method is followed for distinct multi-targeting drug prediction tasks, being independent from the biological questions we intend to address. The only inputs required are a list of targets and a list of anti-targets and the proposed compounds are always selected using nearly the same criteria (in challenge 2 tPSA < 75 was an additional constraint taken into account when selecting compounds).

The solutions presented: For all the problems, the solutions presented are ranked according to what is described in the approach.

Problem 1:

- Solution 1:

ZINC ID: ZINC000040900273

VENDOR ID: Z854859324, MCULE-1723066860

SMILES string: Cc1ccc(NC(=O)c2cccc(C(F)(F)F)c2)cc1NC(=O)c1ccncc1

VENDOR NAME: EnamineStore, Mcule Make-on-demand

Explanation of chemical novelty: Since it has only been presented in one paper, this could be an interesting target to study since the prediction values are good.

- Solution 2:

ZINC ID: ZINC000495355936

VENDOR ID: Z915509806

SMILES string: Cc1ccc(NC(=O)c2ccc3ncccc3c2)cc1C(F)(F)F

VENDOR NAME: EnamineStore

Explanation of chemical novelty: There is no actual presented activities for this compound, so this could be an important time to evaluate this compound.

- Solution 3:

ZINC ID: ZINC000040848081

VENDOR ID: Z854859336

SMILES string: Cc1ccc(NC(=O)c2cccc(C(F)(F)F)c2)cc1NC(=O)c1cccnc1

VENDOR NAME: EnamineStore

Explanation of chemical novelty: The activity of this compound has only been seen on CSF1R gene. By testing this, we could increase the knowledge of the activity of this compound.

- Solution 4:

ZINC ID: ZINC000242823397

VENDOR ID: 183500257, MolPort-043-924-824, 9339742

SMILES string: COc1ccc(NC(=O)c2ccnc(NCCN3CCOCC3)c2)cc1OC

VENDOR NAME: eMolecules, Molport Make-on-demand, Otava (Virtual)

Explanation of chemical novelty: Again, since there is no information about this compound, this could be an opportunity to see if the compound could be useful.

Problem 2:

- Solution 1:

ZINC ID: ZINC000243009863

VENDOR ID: 184290336, MolPort-043-419-559, 9599505

SMILES string: COc1ccccc1N1CCN(c2ccnc(Nc3cccc(Cl)c3)n2)CC1

VENDOR NAME: eMolecules, Molport Make-on-demand, Otava (Virtual)

Explanation of chemical novelty: Again, there is no current knowledge about this compound. With this in mind, this could be an opportunity to evaluate this new compound.

- Solution 2:

ZINC ID: ZINC000243869348

VENDOR ID: 184259625, MolPort-044-173-197, 9589268

SMILES string: CN1CCN(c2ccnc(Nc3ccc(Cl)c(Cl)c3)n2)CC1

VENDOR NAME: eMolecules, Molport Make-on-demand, Otava (Virtual)

Explanation of chemical novelty: The same reason as the previous compound. There are no records for the activity of this compound.

- Solution 3:

ZINC ID: ZINC000084604873

VENDOR ID: 260258169, DA-22711, 260258169, 1341200-40-5

SMILES string: Cc1cnc(Nc2ccc(CN3CCN(C)CC3)cc2)nc1-c1cccc(Cl)c1

VENDOR NAME: eMolecules, Debye Scientific Building Blocks, eMolecules Building Blocks, Hong Kong Chem Here

Explanation of chemical novelty: This compound has a known interaction with AXL, another kinase. This could be important since our targets are also kinases.

**Submission 9662287**

**Authors: Andre Falcao^1^, Samina Kausar^1^**

**Affiliations: ^1^University of Lisbon**

Prediction Methods: The full procedure can be described in three phases, a) a QSAR modeling phase; b) ZINC15 database screening and finally c) Manual curation of the most promising candidates. QSAR MODELING. We set a binary classification QSAR modelling pipeline for all targets and anti-targets of both sub-challenges. We collected data from ChEMBL database. All chemical structures were curated, cleaned and standardized by removing mixtures (handling of unconnected molecules), salt groups, missing data and duplicates. To handle duplicate values for the same molecule the median of activity values was calculated. For this study, only antagonistic Ki, Kd, AC50, EC50, residual activity or activity values were fetched to include data relating to the enzyme inhibition and to ensure that enzyme Binding and inhibition and not binding/inhibition data was considered. After data preparation, all chemical structures were translated using RDKit 2D-descriptors and torsion fingerprints. A combination of which was used to build classification QSAR models for all targets and anti-targets. Positive and negative classes were assigned to all datasets according to the given conditions in the multi-targeting drug DREAM challenge. Moreover, we included all molecules defined by ChEMBL known to be inactive or not active to consider non-binder against all targets (as negatives) and anti-targets (as positives). A state-of-the-art hybrid machine learning procedure was used for the all the models[1]. This procedure couples a RandomForest (RF) based feature selection process was applied to identify and stringently select the most relevant features that are the most adequate for the understudy problems. Before features selection, all datasets were randomly divided into 5-folds to calculate RF based variable importance (VI) from each fold. Thus, features selection approach counts variable importance by calculating the average Mean Decrease Accuracy (MDA) and Mean Decrease Gini (MDG) provided by RF from a series of runs as a tool to rank the predictors. Finally, datasets were again randomly split into training and test sets and the RF VI-based ranked variables were feed to Support Vectors Machines to build the stepwise predictive models and to find a better balance between the biologically relevant set of features and model prediction. An optimised number of features that performed best (maximum predictive performance) across all 5-folds was selected as set of most relevant features. Each model's performance was assessed using the Matthews Coefficient Correlation (MCC). Final QSAR models for all problems were generated using the whole training data with the selected optimized number of features. ZINC15 DATABASE SCREENING. For the database screening all 13,088,593 molecules from ZINC15 database were downloaded applying the filters as purchasable and in-stock molecules. The same RDKit 2D-descriptors and torsion fingerprints were calculated for all molecules and predictions were made from the generated QSAR models for Challenge Questions 1 and 2 for both targets and anti-targets. All screened molecules were ranked using Challenge Question 1 and 2 scoring schemes. In this way, 1325 top-ranked molecules for Challenge Question 1 and 12031 top ranked molecules for Challenge Question 2. Since, model predictability is considered more reliable if the external test molecules fall within the applicability domain (AD) of the predictive models. To analyse the AD of the generated models, Euclidean distances based on the selected number of features (descriptors and torsion fingerprints) for each model were used among all training compounds of corresponding models and the zinc15 compounds. Moreover, the Tanimoto coefficient (ECFP6) was calculated for top-ranked set of compounds to their nearest neighbours in the training set of all targets and anti-targets so to identify the most reliable predictions. All candidate molecules were re-ranked by removing the molecules whose nearest neighbours were from negative classes. After this stringent screening, only 43 molecules were left for Challenge Question 1 and 989 molecules for Challenge Question 2. To eliminate very similar candidate molecules, these were clustered together making sure that each cluster center was far from all the other clusters at least 0.5 using ECFP6 fingerprints. From the top clusters with more consistent and highest predictions, the more promising candidate, the one with the highest score) was selected. MANUAL CURATION. A final manual curation of the best candidate molecules was performed by searching ZINC referenced suppliers for each molecule to ensure that the purchasability of each candidate is within the specifications of the challenge. Also each structure was further searched in SureChembl to verify whether each candidate was not the subject of a previous patent in any of the targets or anti-targets. REFERENCE. Kausar, S. & Falcao, A. O. An automated framework for QSAR model building. J. Cheminform. 10, 1 (2018).

Rationale:

Why is your approach innovative?: There are several innovative issues to the way this screening was conducted. a) the Random Forest Feature Selection procedure, despite having been published by this team, is far from main stream, as well as the automated procedure where the full hybrid methodology has been developed; b) The successsive refinement of the database screening is, to our knowledge, new. It couples the selection of molecules withion the applicability domain to a stringent selection of different candidates through a scalable clustering procedue that ensures that only sufficinetly different candidate molecules are submitted.

Why will your approach be generalizable?: With the exception of the final manual curation procedure which operates on the very best candidates, there is very small manual intervention in the full procedure. In effect after the training datasets were produced for each target of both challenges, the same procedure was autometed and applied to both. Therefore we strongly believe this procedure is generalizable.

Problem 1:

- Solution 1:

ZINC ID: ZINC000008718698

VENDOR ID: MCULE-9456990225

SMILES string: COc1ccc(Br)cc1CN(C)CC(=O)Nc1c(C)n(C)n(-c2ccccc2)c1=O

VENDOR NAME: Mcule

Explanation of chemical novelty: There are no experimentally validated annotation of this molecules for our targets and not found in SureChembl.

- Solution 2:

ZINC ID: ZINC000035946120

VENDOR ID: V029-6643

SMILES string: CC(C)NC(=O)N(Cc1cc(NC(=O)c2ccccc2Cl)ccc1N(C)C)CC1CC1

VENDOR NAME: ChemDiv

Explanation of chemical novelty: There are no experimentally validated annotation of this molecules for our targets and not found in SureChembl.

- Solution 3:

ZINC ID: ZINC000100714025

VENDOR ID: R535109|ALDRICH

SMILES string: CCN(CC)c1ccc(/C=N/c2c(C)n(C)n(-c3ccccc3)c2=O)c(O)c1

VENDOR NAME: Aldrich CPR

Explanation of chemical novelty: There are no experimentally validated annotation of this molecules for our targets and not found in SureChembl.

- Solution 4:

ZINC ID: ZINC000072165851

VENDOR ID: MCULE-4033243382

SMILES string: COc1ccc(-n2ccnc2-c2c(C)n(C)n(-c3ccccc3)c2=O)cn1

VENDOR NAME: Mcule

Explanation of chemical novelty: There are no experimentally validated annotation of this molecules for our targets and not found in SureChembl.

- Solution 5:

ZINC ID: ZINC000019626956

VENDOR ID: MCULE-3745753976

SMILES string: COC(=O)COc1ccc(OC)c(NC(=O)c2c(C)n(C)n(-c3ccccc3)c2=O)c1

VENDOR NAME: Mcule

Explanation of chemical novelty: There are no experimentally validated annotation of this molecules for our targets and not found in SureChembl.

Problem 2:

- Solution 1:

ZINC ID: ZINC000010213695

VENDOR ID: MCULE-5526562762MCULE-9647925319

SMILES string: Cc1cccc(Nc2nc(Nc3ccc(C)c(C)c3)nc(N3CCCC3)n2)c1

VENDOR NAME: Mcule

Explanation of chemical novelty: There are no experimentally validated annotation of this molecules for our targets and not found in SureChembl.

- Solution 2:

ZINC ID: ZINC000019359000

VENDOR ID: MCULE-3630729024MCULE-4308257206

SMILES string: COc1cc(CN2CCN(Cc3cccc4ccccc34)CC2)cc(OC)c1O

VENDOR NAME: Mcule

Explanation of chemical novelty: There are no experimentally validated annotation of this molecules for our targets and not found in SureChembl.

- Solution 3:

ZINC ID: ZINC000019366532

VENDOR ID: MCULE-1506743248

SMILES string: Oc1ccc2ccccc2c1CN1CCN(Cc2c(O)ccc3ccccc23)CC1

VENDOR NAME: Mcule

Explanation of chemical novelty: There are no experimentally validated annotation of this molecules for our targets and not found in SureChembl.

- Solution 4:

ZINC ID: ZINC000041005045

VENDOR ID: MCULE-7312887920

SMILES string: CCOc1ccc2ccccc2c1-c1nc2ccccc2[nH]1

VENDOR NAME: Mcule

Explanation of chemical novelty: There are no experimentally validated annotation of this molecules for our targets and not found in SureChembl.

- Solution 5:

ZINC ID: ZINC000089794014

VENDOR ID: MCULE-7481034537

SMILES string: O=C(Nc1ccc2c(ccn2CCN2CCOCC2)c1)N1CCN(Cc2cccnc2)CC1

VENDOR NAME: Mcule

Explanation of chemical novelty: There are no experimentally validated annotation of this molecules for our targets and not found in SureChembl.

**Submission 9662285**

**Authors: Gregory Koytiger^1^**

**Affiliations: ^1^Immuneering Corporation**

Prediction Methods: Our model, which we term [ModelNameHidden], is a combination convolutional-recurrent neural network that predicts the affinity of drug-protein binding using a numerical representation of the molecule via a variational autoencoder framework (R Gómez-Bombarelli et al, 2016) and the amino acid sequence of the protein. [ModelNameHidden] predicts binding of the molecule at each position of the protein sequence and outputs a confidence score for that positional prediction. The overall predicted affinity is calculated from the weighted average (dot product) of the affinity and confidence score. Using this approach, our model is able to infer the binding site of the molecule in addition to its overall affinity for the protein. The protein sequence is featurized first by splitting it into trimers, yielding three different sequences in different frames. We pre-learn an embedding of the protein trimers using the FastText algorithm run over the entire SwissProt database. These trimer embeddings are then fed into a convolutional layer of size 3, which is then fed into a dilated convolution layer of size 3 and dilation 3, which then is fed into another dilated convolution layer of size 3 and dilation 9. These three convolutional layers are then concatenated along with the original trimer embedding to create a multi-scale representation of each protein position. We also concatenate Pfam domain annotations of each sequence position in an embedding layer created by converting the numeric representation of each domain into a dense vector. Further, we then concatenate the molecule representation by tiling across the full length of the protein sequence. The resultant representation is fed into a Bidirectional Long Term- Short Term (LSTM) recurrent neural network which converts these features into the positional binding and confidence predictions. A final dot product layer summarizes the normalized confidence predictions with the binding positional binding affinity. This approach allows us to use the entire ChEMBL 23 dataset with over 2 million binding affinities to learn one unified binding model that can take as input any protein sequence and SMILES string. We are even able to learn the effect of mutations on binding by feeding in sequence variants as annotated by ChEMBL. We are happy to provide more graphical representations of our model to help visualize the overall architecture. We are also happy to provide the predicted binding sites for the DREAM challenge predictions.

Rationale:

Why is your approach innovative?: The most novel aspect of our model is that we do not only make one prediction for the overall binding but additionally predict along the entire sequence of the protein. This helps the model to learn an explanatory, mechanistic understanding of binding from sequence features. We also use a hybrid dilated convolutional - LSTM neural network that is better able to learn deep sequence features that are predictive of binding to a given molecular architecture. Our use of an embedded domain representation is also innovative and better helps our model identify the correct binding mechanism. These innovative features all stem from the fundamental design principle of [ModelNameHidden], that binding is not a property of a protein but a property of sub-sequence of the protein. This approach to the machine learning prediction of binding is novel because it avoids the creation of a collapsed (and thereby highly lossy) representation of the protein. We also predict affinity instead of binding as a binary output which is pretty difficult and quite novel for a machine learning model.

Why will your approach be generalizable?: The first aspect that makes our model generalizable is that the inputs to our model are simply a sequence representation of a protein and a SMILES string, allowing our single model to predict any desired interaction. Our approach also generalizes well because of the sheer amount of data that is fed into our model. We do not train one model per protein but instead use all known binding interactions to learn the general properties of binding. We further use all of SwissProt to learn effective representations of amino acid sequences through FastText algorithm. This allows us to learn sequence similarity of different amino acid trimers without using any binding data. We also use PFAM to annotate domains which are responsible for a significant portion of binding. The molecular representation of the drug is also similarly learned in an unsupervised way from the entirety of ChEMBL through a variational autoencoder. Thus, even before the model is shown any binding data it already has significant amount of prior information that helps it isolate the correct mechanism. We further show this generalizability by splitting our training and testing data by protein instead of by molecule. We evaluated multiple deep learning architectures and the one that we present here is the only one that is capable of performing well in this test. When we correlate experimental replicate log IC50s for the test proteins we observe an 0.80 Pearson rho. Averaging these replicates and correlating them to the model predicted log IC50, we see a 0.76 Pearson rho. This result shows that even when the model has not seen a single binding interaction for a given protein, it is still able to achieve near experimental accuracy for the prediction. Further the model can be shown to be generalizable because it accurately identifies the residues that are responsible for binding, showing that it converges to the true mechanism - even though no mechanistic ground-truth was ever given to the model. For instance, in our prediction of Gleevec binding to ABL1, we can plot the predicted binding affinity along the sequence and show that our model restricts its predictions overwhelmingly to the protein kinase domain of ABL1 and achieves maximal prediction near the 'gatekeeper' Thr315 residue which is considered among the most important for Gleevec binding. Furthermore, because our model learns binding using a multi-scale sequence representation, it will be able to borrow information effectively across proteins to make truly novel and generalizable predictions.

Problem 1:

- Solution 1:

ZINC ID: ZINC02093883

VENDOR ID: MCULE-9576079569

SMILES string: CC(C)OCCCN1CC(=O)N2[C@H](C1=O)Cc3c4ccccc4[nH]c3[C@@H]2c5ccccc5

VENDOR NAME: Mcule

Explanation of chemical novelty: Visibly different, lacks the quinazoline core common to many TKIs

- Solution 2:

ZINC ID: ZINC02833706

VENDOR ID: 7472894

SMILES string: c1ccc(c(c1)C(=O)[O-])NC(=S)NC(=O)/C=C/c2cccc(c2)[N+](=O)[O-]

VENDOR NAME: ChemBridge

Explanation of chemical novelty: Very distinctive from current TKIs

Problem 2:

- Solution 1:

ZINC ID: ZINC84724044

VENDOR ID: M-124001

SMILES string: C[C@@H]1CC[C@H]2C(=C1)C=C[C@@H]([C@]23CCC(=O)O3)C

VENDOR NAME: TLC Pharmaceutical Standards

Explanation of chemical novelty: Distinctive from traditional kinase inhibitors

- Solution 2:

ZINC ID: ZINC00001266

VENDOR ID: Y0427

SMILES string: C[C@@H](C1=[NH+]CCN1)Oc2c(cccc2Cl)Cl

VENDOR NAME: AK Scientific

Explanation of chemical novelty: While the compound, lofexidine, is itself well known, the use in this context would be novel

**Submission 9662215**

**Authors: Joerg Kurt Wegner^1^, Huub Henkelsma, Gerard JP van Westen^2^, Brandon Bongers^3^, Lindsey Burggraaff^3^, Jesper Van Engelen^3^, Xuhan Liu^3^, Xuhan Liu, Marina Gorostiola Gonzalez, Marvin Steijaert Hugo Gutiérrez de Teran^4^, Holger Hoos, Anthe Janssen**

**Affiliations: ^1^Janssen Pharmaceuticals, ^2^Leiden Academic Center for Drug Research, ^3^Leiden University, ^4^Uppsala University**

Prediction Methods: Our methods are described in our project wiki and on google docs, https://docs.google.com/document/d/1JEWeV99FDk1zwV5Hu4tRUYgSeJHPFpzqFj9MOZl3sU0/edit?usp=sharing . Solutions (compounds) are ranked from 1 to 5. Compounds have been checked for novelty and patentability.

Rationale:

Why is your approach innovative?: We are confident that our approach represents one of the most rigorous methods of all submissions. 1) We tailored our workflow to every target and anti target, using benchmark data. 2) This was done for multiple models (statistical, structure based, and metadynamics). 3) Statistical models (biosignatures) trained on Janssen data were used, in general these models achieve a really predictive ROC compared with statistical models trained on public data (e.g. ROC ~0.95 compared with ROC ~0.8). 4) Metadynamics simulations gave us more confidence that the binding modes of compounds that we selected were stable.

Why will your approach be generalizable?: In our lab we have used and are using comparable workflows for polypharmacology predictions. We have applied these workflows on different target classes such as enzymes and transmembrane receptors. By using large libraries containing compound and target information, our approach is generalizable to proteins with sufficient bioactvity data.

Problem 1:

- Solution 1:

ZINC ID: ZINC000012493340

VENDOR ID: ASN02751147,ST50530067,17089330,MCULE-8541271424

SMILES string: COc1ccc2cc(CNc3ccc(Cl)cc3)c(=O)[nH]c2c1

VENDOR NAME: Asinex,TimTec,eMolecules,Mcule

Explanation of chemical novelty: This compound was selected because it was one of the top scoring compounds in all of the models, the binding mode looked good in all on-targets, and it scored relatively well in the Janssen statistical model. Moreover this scaffold (1H-quinolin-2-one) would represent a new scaffold for RET. Additionally, this compound is not similar to known active compounds for the challenge proteins and not present in SciFinder.

- Solution 2:

ZINC ID: ZINC1801746

VENDOR ID: R853313|ALDRICH

SMILES string: Clc1ccc(Oc2ncnc3scc(-c4ccc(Br)cc4)c23)c(Cl)c1

VENDOR NAME: Aldrich CPR (there are more than 5 vendors)

Explanation of chemical novelty: This compound scored really well in the metadynamics, had a good binding mode in the on-targets, but a lower score in the statistical models. This compound is not similar to known active compounds for the challenge proteins and not present in SciFinder.

- Solution 3:

ZINC ID: ZINC000001398621

VENDOR ID: 7N-768

SMILES string: Nc1nc(Nc2ccc(Cl)cc2)sc1C(=O)c1ccc(F)cc1

VENDOR NAME: KeyOrganics (there are more than 5 vendors)

Explanation of chemical novelty: The binding mode of this compound looks really good in RET (triple interaction with the hinge), however it was not able to dock well in BRAF. This compound is not similar to known active compounds for the challenge proteins and not present in SciFinder.

- Solution 4:

ZINC ID: ZINC000004032944

VENDOR ID: MolPort-003-033-441, F0611-0801, 4937996, MCULE-8382667119

SMILES string: Cc1ccc2cc(CCNC(=O)c3ccc(Cl)cc3Cl)c(=O)[nH]c2c1C

VENDOR NAME: Molport SC Economical, Life Chemicals, eMolecules, Mcule

Explanation of chemical novelty: This compound is similar to Solution 1, it scored better in the structure based models, but worse in the Janssen statistical model. This compound is not similar to known active compounds for the challenge proteins and not present in SciFinder.

- Solution 5:

ZINC ID: ZINC000095930125

VENDOR ID: HY-15730

SMILES string: C=CC(=O)N1CCC(Oc2cc3c(cc2OC)ncnc3Nc2ccc(Cl)c(Cl)c2F)CC1

VENDOR NAME: MedChem Express Economical (there are more than 5 vendors)

Explanation of chemical novelty: Although this compound, Poziotinib, is not novel it might be interesting to test from a drug repurposing perspective, no data is given in ChEMBL for this compound.

Problem 2:

- Solution 1:

ZINC ID: ZINC000020351955

VENDOR ID: 77807007, MolPort-005-135-808, 23830893, MCULE-7651088158

SMILES string: O=C(c1ccc2nccn2c1)N1CCc2[nH]nc(CCC3CCCC3)c2C1

VENDOR NAME: ChemBridge Economical, Molport SC Economical, eMolecules, Mcule

Explanation of chemical novelty: This compound has a good activity profile in the structure-based models. Additionally, this compound scored good in the Janssen statistical model. This compound is not similar to known active compounds for the challenge proteins and not present in SciFinder.

- Solution 2:

ZINC ID: ZINC000067817173

VENDOR ID: 45568190, MolPort-019-889-395, 36585474, MCULE-5856441480

SMILES string: Cc1nc(C)c2c(n1)O[C@H](CN1CCc3[nH]nc(-c4ccccc4F)c3C1)C2

VENDOR NAME: ChemBridge Economical, Molport SC Economical, eMolecules, Mcule

Explanation of chemical novelty: This compound has a good bioactivity profile in structure-based models, except for PIK3CA. However, PIK3CA inactivity is predicted more favorably by statistical models. This compound is not similar to known active compounds for the challenge proteins and not present in SciFinder.

- Solution 3:

ZINC ID: ZINC000015729174

VENDOR ID: MolPort-007-787-103, 16249573, MCULE-9202236133

SMILES string: COc1ccc(-c2[nH]ncc2CN2CCN(C(=O)Cc3ccc(F)cc3)CC2)cc1

VENDOR NAME: Molport SC Economical, eMolecules, Mcule

Explanation of chemical novelty: The binding poses of this compound in AURKA and PAK1 looked good. Also, it scored good in metadynamics. This compound scored good in the Janssen statistical model. Additionally it has a good binding pose in AURKA and PAK1. This compound is not similar to known active compounds for the challenge proteins and not present in SciFinder.

- Solution 4:

ZINC ID: ZINC000257201200

VENDOR ID: BDE30723429, 43249057

SMILES string: Cn1cc(CN2CCC(c3ccnc(Nc4cnccn4)c3)CC2)cn1

VENDOR NAME: Asinex, eMolecules

Explanation of chemical novelty: This compound scored relatively good in the Janssen statistical model. This compound is not similar to known active compounds for the challenge proteins and not present in SciFinder.

- Solution 5:

ZINC ID: ZINC000035509152

VENDOR ID: MolPort-010-686-472

SMILES string: O=C(CCc1ccccc1)NCCc1n[nH]c(-c2ccccc2)n1

VENDOR NAME: Molport SC Economical (there are more than 5 vendors)

Explanation of chemical novelty: The binding poses of this compound in the required on-targets looked good. Additionally, TAK1 binding seems unlikely. This compound is not similar to known active compounds for the challenge proteins and not present in SciFinder.

**Additional Methods Information from team SuperModels:**

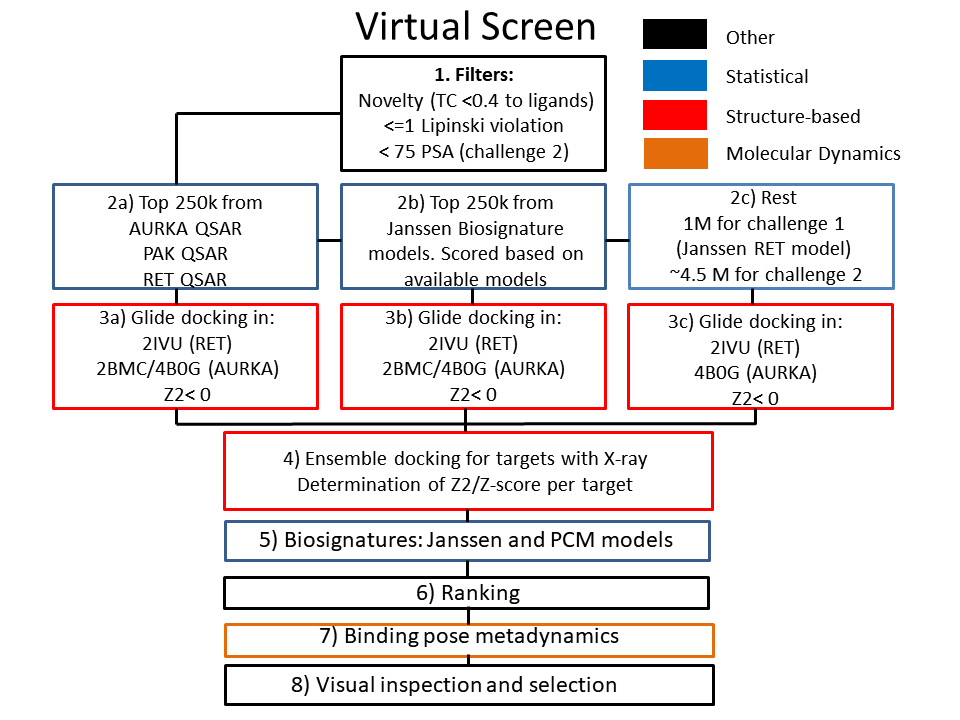

*Figure 1: Overview of the used workflow. The different stages and filters are further discussed below.*

**1. Filtering of ZINC**

ZINC-15^1^ as accessed on January 8th was processed in BioVia Pipeline Pilot 2016 (box 1). The following criteria as proposed by the challenge were used as hard filter: maximum 1 Lipinski violation, Novelty (ECFP_6, TC <0.4 to known ligands in ChEMBL23^2^), and a PSA below 75 for challenge 2.

**2a & 5. Statistical models based on public data**

We created models based on public data from the following sources: ChEMBL23^2^, data from Klaeger *et. Al,^3^* Eidogen^4^ and from ExCAPE-DB^5^. The training dataset was filtered to contain only compounds tested on the challenge kinases for use in QSAR and additional related kinases for PCM. The compounds were filtered on a molecular weight lower than 700. FCFP4 and physical chemical descriptors were generated for QSAR modeling. Additional protein descriptors were calculated based on full sequence alignment for PCM modeling. We created QSAR models for RET, AURKA and PAK1 (box 2a). These models were used as initial screen for docking a subset of the ZINC set (box 2c). PCM models were created as described in a previous study.^6^ These PCM models were applied in scoring the ZINC compounds in box 5 “biosignatures” for targets for which no Janssen model was available. The statistical models were optimized by tuning the random forest model hyperparameters using random search, a basic approach to automated machine learning.^7^ Further details of used model parameters, descriptors and preliminary results are shown in the Supplementary methods and results.^8^

**2b & 5.** **Statistical modelling - Janssen Pharmaceutica**

Janssen biosignatures were created based on a framework for biosignature data fusion models, which will be released in combination with the corresponding papers.^9, 10^ This method, in short, uses multiple individual logistic regression models trained on the data. The predicted scores of the individual models are turned into probabilities using Platt scaling and are fused using a NoisyOR function. The individual logistic regression models differ in the type of features that are used: atom pair 2D fingerprints, extended connectivity fingerprints (ECFP6), physicochemical descriptors (calculated by MOE), and shape based comparison of a compound against bioactive conformations of RCSB PDB ligands (FastROCS). Further information can be found on github: <https://github.com/bioinf-jku/project_BBDD>. Those models result in probability estimates for compound-gene pairs. These probabilities were then used to predict the ZINC compounds on genes of interest. The probabilities were multiplied by the number of points for a target in the challenge, subtracted for anti-targets and summed up for targets. Those ranks were subsequently used in the ranking of 250k additional compounds (2b). For several of the targets, models were missing due to the unavailability of sufficient data and/or poor performance in validation. For those targets PCM models were constructed as further described above.

**3 & 4. Structure-based**

Structure based modelling was performed using the Schrödinger 2017-4 suite^11^, and using the OPLS3 force field^12^. ZINC compounds were prepared using Ligprep. Docking and ligand preparation of the “rest” subsets (2c, figure 1) was done using ligprep and glide using the Schrödinger 2017-2 suite, which was the available version at the Stallo supercomputing center at The Arctic University of Norway (UiT). In retrospective experiments this version achieved comparable enrichments (see Supplementary methods and results). Due to time constraints, for challenge 1 we docked the remaining top 1M based on the Janssen model for RET. For challenge 2 all compounds were docked. For every target with available X-rays we rigorously benchmarked which workflow to use. For the benchmarking we created Active/Inactive/Decoy (AID) sets using the DUD-e webserver.^13^ Subsequently all available X-rays with a co-crystallized ligand were downloaded and prepared using the protein preparation wizard. Enrichments were calculated (BEDROC (alpha =160.9) and ROC), and for a subset of structures we calculated SPLIF scores^14^. On the basis of these results we normalized (Z-score) the SPLIF and docking scores with respect to the actives from the AID-set for that target. By doing so we optimized the workflow to calculate an ensemble, Z2 score^15^, this Z2 score increased the enrichment for 10 out of the 13 targets. A more detailed methods section and preliminary results can be found in the supplementary methods and results section.^8^

**6. Ranking**

The ranking prior to metadynamics was based on the sum of the individual ranks of targets and antitargets (inverse ranking). The weights per target were based on the number of points per target (1, 3 or 5) and the performance of statistical/structure based models in retrospective experiments. Herein the ROC was used for Janssen models, and the ROC weighted by the number of samples for the PCM models. For the structure based results we used the sum of the BEDROC (alpha =160.9) and ROC. Targets for which induced fit docking was used to generate more protein structures were penalized by multiplying with 0.5. Targets with less than 100 actives in the benchmark set were penalized by multiplying with the fraction of number of actives. The final weighting scheme is provided in the Supplementary methods and results.^8^

**7. Binding Pose metadynamics**

For a subset of ligands we calculated a Metadynamics-composition score^16^, this score was added to the existing docking scores. In retrospective tests this increased the reranking of compounds (top100) for both RET and AURKA but not for PAK1 (see Supplementary methods and results).^8^ We used the following PDB codes: 2IVU for RET and 2BMC for AURKA. All termini were capped. The Metadynamics-composition scores were used to rerank the top *n* ligands for those targets and used in the further decision making.

**8. Visual inspection and final selection**

Visual inspection was performed on the majority of well scoring compounds by the majority of our team. Compounds were inspected based on their 2D structure (i.e. hinge binder), patentability, and 3D docking pose on the required targets. Patentability was checked in both Surechembl^17^ and the IBM patent system.

14.	Da, C.; Kireev, D., Structural protein–ligand interaction fingerprints (SPLIF) for structure-based virtual screening: method and benchmark study. J. Chem. Inf. Model.

**2014**, 54, 2555-2561.

15.	Sastry, G. M.; Inakollu, V. S.; al., e., Boosting virtual screening enrichments with data fusion: coalescing hits from two-dimensional fingerprints, shape, and docking. *J. Chem. Inf. Model.* **2013**, 53, 1531-1542.

16	A.J. Clark; P. Tiwary; al., e., Prediction of protein–ligand binding poses via a combination of induced fit docking and metadynamics simulations. *J. Chem. Theory Comput.* **2016**, 12, 2990-2998.

17	Papadatos, G.; Davies, M.; Dedman, N.; Chambers, J.; Gaulton, A.; Siddle, J.; Koks, R.; Irvine, S. A.; Pettersson, J.; Goncharoff, N., SureChEMBL: a large-scale, chemically annotated patent document database. Nucleic acids research **2015**, 44, D1220-D1228.

**Submission 9662213**

**Authors: Zhaoping Xiong^1^**

**Affiliations: ^1^ShanghaiTech University**

Prediction Methods: We built a triple classifier to predict the bioactivity of a compound to a target protein.

The classifier encodes both ligand molecules and target proteins by constructing a new end-to-end differentiable

neural net architecture, in which proteins are represented by amino acid sequence embedding and molecules

by fingerprint generated from graph convolutional neural network.

Rationale:

Why is your approach innovative?: Graph convolutional neural network for molecular fingerprint is often used

as a deep learning model to predict the bioactive. As a ligand-based model, the inability to predict

bioactivity to a new target with few bioactive compounds constrains its generalization.

To build a more generalizable and useful deep model, we incorporate the feature from target proteins

by converting their amino acid sequence into “word embedding” vector and trained together with

molecular fingerprint generated from graph convolutional neural network.

Why will your approach be generalizable?: For a given new compound-target pair, our predictive model will classify

the bioactivity into potent, weak or inactive. Our deep model is trained on a high-quality dataset

and learns features from both ligands and targets, achieving a predictive accuracy of 0.91 on a external test set.

Problem 1:

- Solution 1:

ZINC ID: ZINC98209221

VENDOR ID: 5429

SMILES string: CCN1CCN(Cc2ccc(NC(=O)c3ccc(C)c(Oc4ccnc5[nH]ccc45)c3)cc2C(F)(F)F)CC1

VENDOR NAME: Tocris

Explanation of chemical novelty: It has few citations and returns only 46 compounds by similarity search with a threshold of 0.8 on scifinder.

Only patented as inhibitor of MAP4K2 and reported to inhibit TAK1, LYN and Abl kinase.

Problem 2:

- Solution 1:

ZINC ID: ZINC147474927

VENDOR ID: F20815

SMILES string: COc1ccc(N(C(=O)Nc2c(C)cccc2C)c2cc(Nc3ccc(N4CCN(C)CC4)cc3)ncn2)c(OC)c1

VENDOR NAME: AstaTech

Explanation of chemical novelty: Returns only 42 compounds by similarity search with a threshold of 0.8 on scifinder.

- Solution 2:

ZINC ID: ZINC3938668

VENDOR ID: 2693

SMILES string: Cc1[nH]c(/C=C2\C(=O)Nc3ccc(S(=O)(=O)Cc4c(Cl)cccc4Cl)cc32)c(C)c1C(=O)N1CCC[C@@H]1CN1CCCC1

VENDOR NAME: Tocris

Explanation of chemical novelty: Potent, selective and ATP-competitive inhibitor of MET kinase (IC50 values are 9, 68, 200,

1400, 3000, 3800 and 6000 nM for MET, Ron, Flk-1, c-abl, FGFR1, EGFR and c-src respectively and > 10000 nM for IGF-IR,

PDGFR, AURORA2, PKA, PKBα, p38α, MK2 and MK3).

Additional detailed writeup from Zhaoping Xiong:

Fusing protein embedding with convolutional neural fingerprints for bioactive prediction

Zhaoping Xiong^1,2,3^, Mingyue Zheng^*,2^ , Hualiang Jiang^*,1,2^

*^1^School of Life Science and Technology, ShanghaiTech University, Shanghai 200031, China;*

*^2^Drug Discovery and Design Center, State Key Laboratory of Drug Research, Shanghai Institute of Materia Medica, Chinese Academy of Sciences, 555 Zuchongzhi Road, Shanghai, 201203, China;*

*^3^University of Chinese Academy of Sciences, No.19A Yuquan Road, Beijing 100049, China*

### **Results**

#### **Performance on proof of concept tests**

Table 1 | Predictive performance on regression models

| **Regression models**  **(RMSE of pIC50)** | **Train (80%)** | **Valid (10%)** | **Test (10%)** |
| --- | --- | --- | --- |
| Predict as the mean of the target | 0.605 | 0.582 | 0.594 |
| ECFP (multi-task) | 0.486 | 0.521 | 0.525 |
| Neural Fingerprints (multi-task) | 0.458 | 0.506 | 0.508 |
| ECFP + Protein Embedding | 0.421 | 0.477 | 0.480 |
| Neural Fingerprints + Protein Embedding | **0.294** | **0.417** | **0.414** |

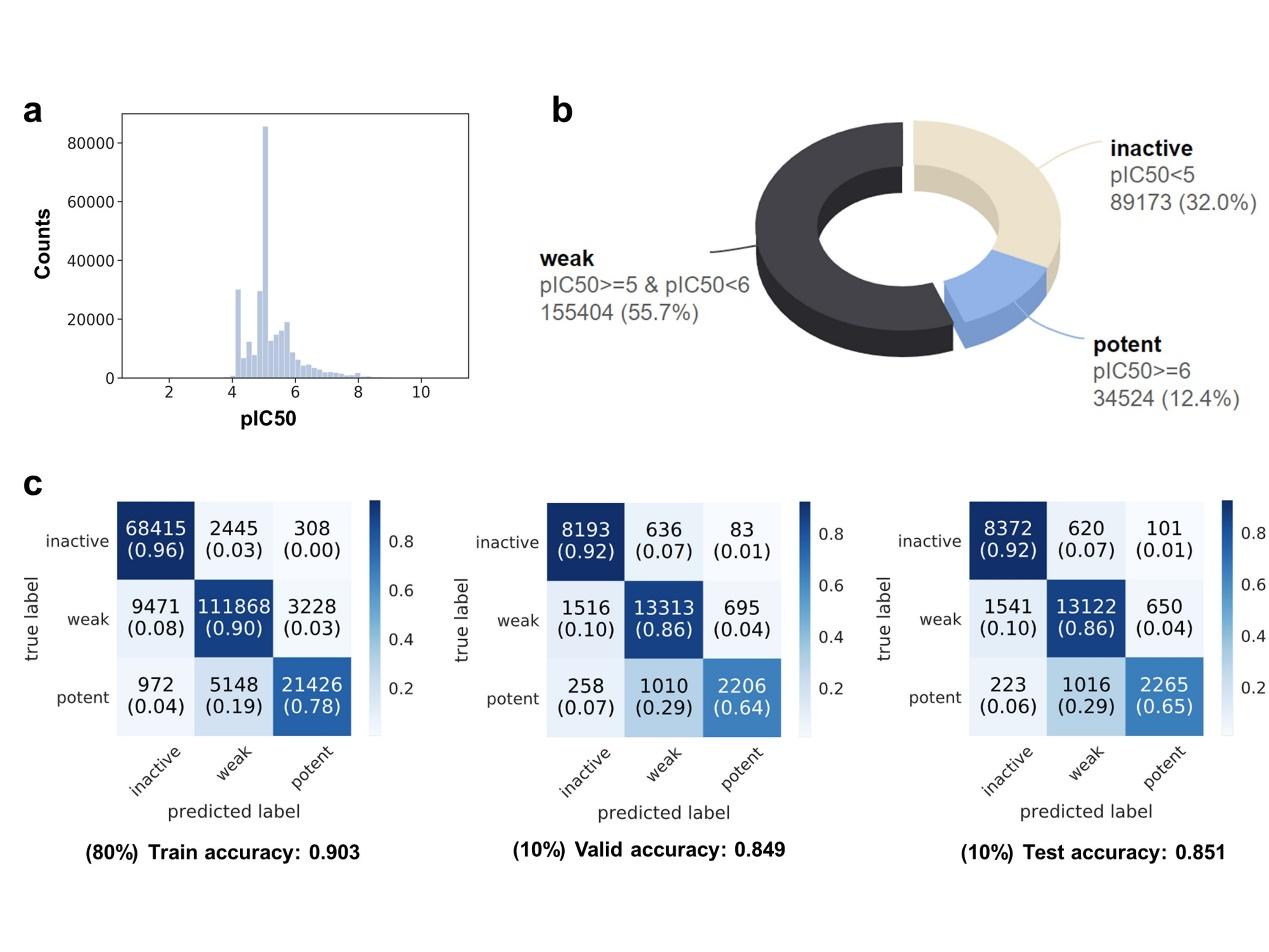

Fig. 1 | Recapitulating the final three-class classification model used for screening. a, The distribution of pIC50; b, The distribution of classes; c, The accuracy of the model across train, valid and test sets.

In order to set the baseline, we firstly built regression models and compare the performances of different models. As shown in Table 1, ligand-based methods, such as multi-task neural networks that take as input ECFP or Neural Fingerprints, will have some prediction power over random guess the value with mean. When protein embeddings are also included for modeling, the overall predictive performance increased noticeably. Also, Neural Fingerprints performs better than ECFP. Fig. 1a shows us that the pIC50 data points are in deformed distribution that concentrates extremely on center, which makes the model tend to predict as those majority mediocre values to reduce loss for most samples when training, resulting in non-differentiable models to values in the two ends. But the values in the two ends are more our concern. Therefore, as in Fig. 1b, we classify the bioactivity as inactive, weak and potent with criteria of pIC50<5, pIC50>=5 & pIC50<6 and pIC50>=6, respectively. And then, a three-class classification model is built and Fig. 1c reports the the predictive performance. We see potent class has high tendency to be misclassified as weak class, which will later be considered when selecting compounds for the problems we would like to validate with experiments.

### **Datasets**

The bioactive data used to build fused embedding model are curated by Merget and Fulle et.al. and taken from https://github.com/Team-SKI/Publications^1^, which collects data from four different sources—the Tang set^2^ (a collection of the kinase profiling data sets of Metz^3^, Davis^4^ and Anastassiadis^5^), PKIS^6–8^, Christmann2016^9^ and a curated ChEMBL kinase inhibitor panel by Merget^10^. We merged the measurements from different sources by mean, kept compounds with more than 20 bioactive data and kinases with more than 20 compounds, resulting in 392 kinases, 2140 compound and 279101 data points of pIC50 (~33.3% coverage).

To facilitate later validation, we constructed a virtual screening library from the commercially available compound libraries, which incorporates the company Selleckchem’s five compound libraries, including the FDA-approved Drug Library, Preclinical and Clinical Compound Library, Bioactive Compound Library-I, Kinase Inhibitor Library and Natural Product Library (Table 2).

Table 2 | The virtual screening library.

| **Library id** | **Library name** | **Volume** |
| --- | --- | --- |
| L1300 | FDA-approved Drug Library | 2557 Compounds |
| L3900 | Preclinical and Clinical Compound Library | 2542 Compounds |
| L1700 | Bioactive Compound Library-I | 5338 Compounds |
| L1200 | Kinase Inhibitor Library | 766 Compounds |
| L1400 | Natural Product Library | 2113 Compounds |

The protein sequence embedding model is pre-trained on all kinase related proteins by search with key word ‘kinase’ on UniProt (<https://www.uniprot.org>), including 19741 reviewed kinases sequences at the length of 100-5000. Then the kinase domain sequence of 392 kinases that have bioactivity data in our dataset are extracted to represent those kinases, which will be used to infer the protein embedding of those kinases. All the data and the preprocessing procedure are available at <https://github.com/xiongzhp/Multi-Targeting> (private yet) and can be easily reproduced on Jupyter notebook server.

### **Methods**

#### **Protein sequence embedding**

We intended to incorporate protein information for bioactive prediction. Conventionally, 3D protein structures whether modelled homologically or resolved by crystallography, cryoEM and NMR are used for analyzing structure-activity relationships. However, the infeasibility of high quality structures, solid homology modelling and trustworthy docking scoring methods restrained the usage of protein structure for bioactive prediction on a large scale. So we hope to incorporate protein information directly from the protein sequence by learning an embedded representation like processing natural language, in which protein kinases are differentiated by its sequence, just like different text are represented with different vectors and text with similar content will have similar vectors (measured by distance or cosine similarity).

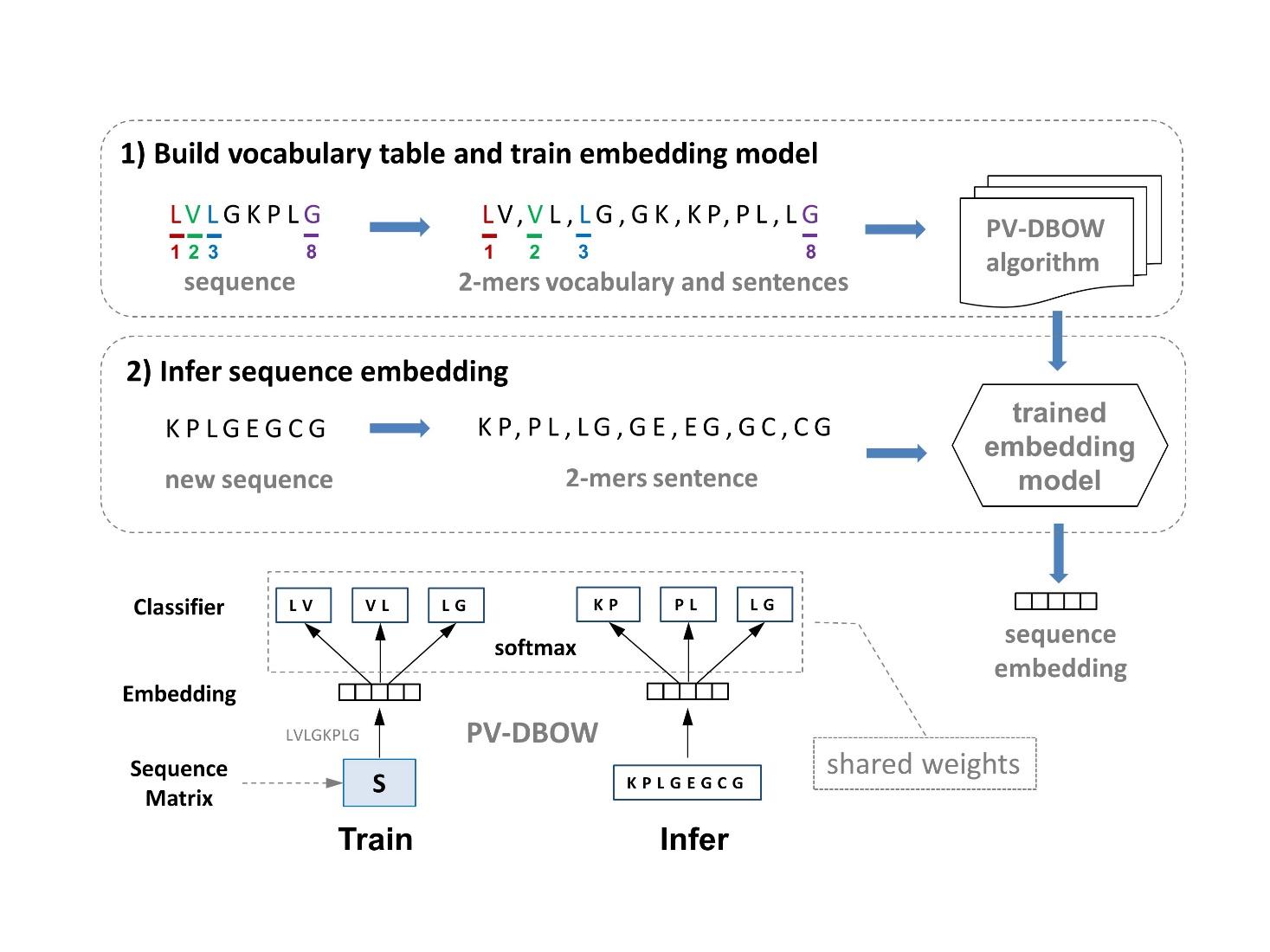

Fig. 1 | Protein embedding scheme. 1) The protein sequences are transformed into ‘sentences’ composed of 2-mers and train the model with PV-DBOW (Distributed Bag of Words version of Paragraph Vector) algorithm, in which the sequence embedding is fed into softmax layer, forming classification tasks for the ‘words’ in a window; 2) Inferring the embedding of a new sequence are based on the trained model by sharing the pre-trained weights in softmax layer.

Learning an embedded representation for sequential data has been well-established in natural language processing and applicable to protein sequences^11–14^. These embeddings can be used to distinguish ordered/disorder proteins or proteins from different families and predict properties, such as localization, T50, absorption and enantioselectivity. However, the protein embedding scheme is usually not as generalizable as in the natural language paradigm. We need to test out which protein embedding scheme works best for the problem of your interest. Fig. 1 demonstrates our final adopted protein embedding scheme, where two consecutive amino acids are combined as the vocabulary of the ‘language’ of protein sequence. So each protein sequence is divided into 2-mers ’words’ and fed into PV-DBOW (Distributed Bag of Words version of Paragraph Vector) model for training. When training with PV-DBOW model, we first sample a window of 2-mers (LV, VL, LG) from a sequence, then the sequence embedding is fed into softmax layer, forming a classification task for the ‘words’ (2-mers) in that window. The sequence embedding updates with stochastic gradient descent when back propagating the cross entropy loss at each iteration. Inferring the embedding of a new sequence are based on the previously trained embedding model by sharing the pre-trained weights in softmax layer.

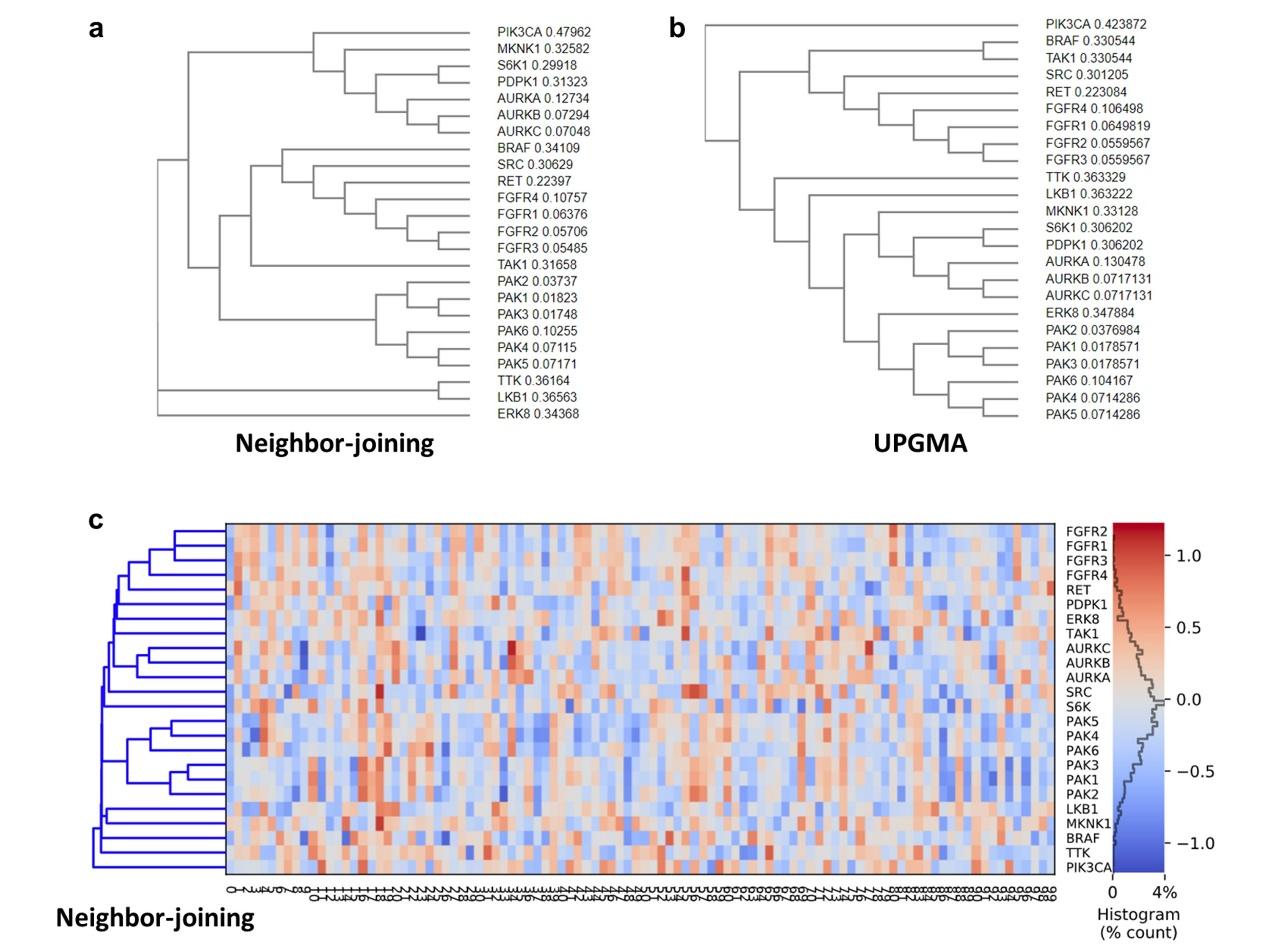

Fig. 2 | Visualization of protein sequence embeddings. The phylogenetic trees of kinase domain sequences are generated from Clustal Omega multiple sequence alignment using different clustering methods, **a,** neighbor-joining and **b,** UPGMA.. **c,** The cluster tree of sequence embeddings mostly agrees with phylogenetic trees, but not exactly the same.

To visualize the protein embedding results, we select the learned embeddings of those pro- or anti- targets in the challenge and some of the proteins in the same subfamily for demonstration. As shown in Fig. 2, the cluster tree of sequence embeddings mostly agrees with phylogenetic trees, but not exactly the same, which implies the deep learning approach perceive the sequence in a slightly different way.

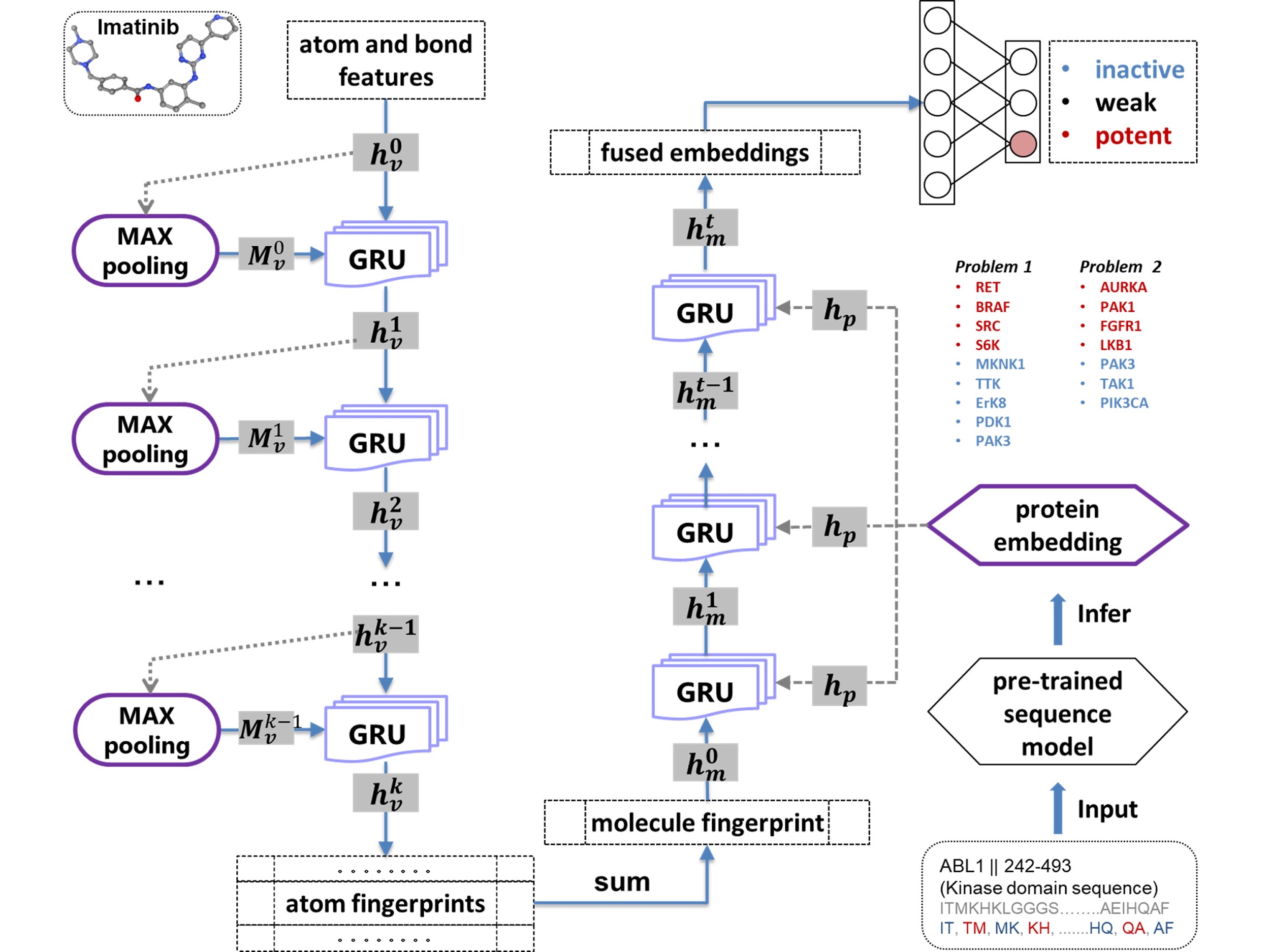

Fig. 3 | The framework of fused embedding. **1)** Given a small molecule, the atoms are featurized into vectors and linear transformed into initial atom state vector $h_{v}^{0}$. For a specific atom $v$, to include information from its environment, the max-pooling is performed on features of neighbor atoms and bonds, obtaining $M_{v}^{0}$**.** GRUs (Gated Recurrent Units) takes as input the state vector $h_{v}^{k-1}$ and $M_{v}^{k-1}$ and outputs new state vector $h_{v}^{k}$ at time step $k$. **2)** $h_{v}^{k}$ are summarized into the molecule ‘fingerprint’ $h_{m}^{t}$ and fed into GRUs together with protein embedding to update state. **3)** The fused embedding $h_{m}^{t}$ (the state vector at time step t) are further fed into a softmax layer for a three-class classification.

Graph convolutional neural network for molecular fingerprint (neural fingerprints) is often used as a deep learning’s approach to predict the molecule properties and bioactivities^15,16^. As a ligand-based model, the inability to predict bioactivity to a new drug target with few bioactive compounds constrains its generalization. To build a more generalizable and useful deep model, we incorporate the feature from the proteins by converting their amino acid sequence into “word embedding” vector (protein embeddings) and trained together with neural fingerprints generated from graph convolutional neural network. There are several studies pre-printed on bioarxiv that predict the affinity of compound-protein binding by incorporating protein embedding with Morgan fingerprints or smiles seq2seq embedding^17,18^. Those performances might be encouraging but not good enough to put into real application with a RMSE of 1.23 for pIC50. Here, we would like to focus on kinase inhibition prediction and model with kinase domain sequences only rather than the whole-length protein sequence used by other studies. Because whole-length protein sequences may induce promiscuous and low differentiation between proteins in the same family, which is unfavorable to our problem-solving. In our fused embedding framework, we mingle protein sequence embedding with neural fingerprints for bioactive prediction. Given a small molecule, the atoms are featurized into vectors and are linear transformed into initial atom state vector $h_{v}^{0}$. Because our neural fingerprint implementation is atom-centric, the bond features are treated as atoms’ neighbor features together with features from neighbor atoms. For a specific atom $v$, in order to include information from its environment, the max-pooling is performed on features of neighbor atoms and bonds, in which the max features across different neighbors are retained, obtaining $M_{v}^{0}$. Each atom will go through several steps of state updating, during which GRUs (Gated Recurrent Units) taking as input the state vector $h_{v}^{k-1}$ and $M_{v}^{k-1}$ and outputting new state vector $h_{v}^{k}$ at time step $k$. 2) The atom ‘fingerprints’ (state vectors at time step $k$) are summarized into the molecule ‘fingerprint’ and fed into GRUs together with protein embedding to update state. This will also iterate several steps (to go deeper), expecting to extract more delicate message. 3) The fused embedding $h_{m}^{t}$ (the state vector at time step t) are further fed into a softmax layer. The cross entropy is used as losses for backpropagation training.

**Submission 9662212**

**Authors: Masahiro Mochizuki^1^**

**Affiliations: ^1^DeNA Co., Ltd.**

Prediction Methods: |

Prediction of Kinases Inhibition with Multi-task Ensemble Learning Model

I obtained a dataset of inhibition assay on 15 enzymes that are targets and antitarget of Problem 1 and 2.

Compounds in ZINC15 were filtered based on criteria below\:

(1) Its purchasability is "in stock".

(2) It satisfies at least 3 of 4 criteria in Lipinski's rule.

(3) Its TPSA was smaller than 75 angstroms squared.

Fingerprints were generated for each compound in the dataset.

Here 8 kinds of fingerprints, namely ECFP, FCFP, Atom Pairs, Topological Torsion, Avalon and RDKit Fingerprint were generated on all compounds.

Regression models of Random Forest were trained with each of fingerprints independently.

Then the models were ensembled by a logistic regression model using stacked generalization.

When a hash function was applied to the raw fingerprint, a technique proposed by Weinberger et al was adopted

in order to solve the questions in multi-task learning paradigm.

As a result, the single ensembled model predicted inhibition activity to all the enzymes.

Expected values of scores were calculated for each compound in the library based on the probability estimates by the model.

Finally, 5 compounds giving the highest expected values of score were submitted to the organizer of this challenge.

Rationale:

Why is your approach innovative?: |

(1) Ensemble of multiple fingerprints.

While multiple learning algorithms are ensembled in usual stacked generalization,

multiple fingerprints are done here. Better predictive performance is expected than each of individual fingerprints.

(2) Hybrid of regression and classification.

The ensembled model is composed of two kinds of learners\: Random Forest regressors and a logistic regression classifier.

Each of the formers accepts a fingerprint as a feature vector, and predicts inhibition rates of each compound.

Then the latter accepts predictions of the formers, and predict whether the inhibition rates are higher than threshold or not.

Hereby the ensembled model is trained with quantitative data, and gives probabilities of binary classification at last.

This mechanism enables to calculate expected values of scores with less information loss caused by binning of inhibition rates.

Why will your approach be generalizable?: |

If training data were provided, my model could be fitted to any targets and off-targets very easily.

In addition, the single model could be generalizable to multiple problems because of multi-task learning.

Actually Problem 1 and 2 were solved using a common model in this challenge.

Problem 1:

- Solution 1:

ZINC ID: ZINC000022200171

VENDOR ID: MolPort-004-757-286

SMILES string: NNc1ccc(Cl)nc1

VENDOR NAME: Molport SC Economical

Explanation of chemical novelty: Maximum Tanimoto similarity to known targets is 0.271.

- Solution 2:

ZINC ID: ZINC000239174455

VENDOR ID: MolPort-001-621-400

SMILES string: O=C1[C@@H]2[C@H]3CC[C@@H](C3)[C@H]2C(=O)N1c1ccc(N2C(=O)[C@H]3[C@H]4CC[C@@H](C4)[C@H]3C2=O)c(Cl)c1

VENDOR NAME: Molport SC Economical

Explanation of chemical novelty: Maximum Tanimoto similarity to known targets is 0.146.

- Solution 3:

ZINC ID: ZINC000000368219

VENDOR ID: MolPort-002-220-470

SMILES string: COc1cc(C)c(/C=C2/NC(=O)NC2=O)cc1C(C)C

VENDOR NAME: Molport SC Economical

Explanation of chemical novelty: Maximum Tanimoto similarity to known targets is 0.217.

- Solution 4:

ZINC ID: ZINC000008729997

VENDOR ID: MolPort-001-757-541

SMILES string: NNc1ccc(Cl)cn1

VENDOR NAME: Molport SC Economical

Explanation of chemical novelty: Maximum Tanimoto similarity to known targets is 0.179.

- Solution 5:

ZINC ID: ZINC000247779175

VENDOR ID: MolPort-002-808-838

SMILES string: O=C1[C@@H]2[C@H]3C=C[C@@H](C3)[C@H]2C(=O)N1c1ccc(N2C(=O)[C@H]3[C@H]4C=C[C@@H](C4)[C@H]3C2=O)c(Cl)c1

VENDOR NAME: Molport SC Economical

Explanation of chemical novelty: Maximum Tanimoto similarity to known targets is 0.146.

Problem 2:

- Solution 1:

ZINC ID: ZINC000004165365

VENDOR ID: MolPort-001-933-900

SMILES string: CC1=C(C(=O)Nc2ccccc2C)[C@@H](c2ccc(Br)cc2)NC(=O)N1

VENDOR NAME: MolPort SC Economical

Explanation of chemical novelty: Maximum Tanimoto similarity to known targets is 0.375.

- Solution 3:

ZINC ID: ZINC000000070848

VENDOR ID: MolPort-001-951-773

SMILES string: CC1=C(C(=O)Nc2ccccc2C)[C@H](c2ccc(F)cc2)NC(=O)N1

VENDOR NAME: MolPort SC Economical

Explanation of chemical novelty: Maximum Tanimoto similarity to known targets is 0.500.

- Solution 4:

ZINC ID: ZINC000005560248

VENDOR ID: MolPort-004-256-659

SMILES string: CC1=C(C(=O)Nc2ccccc2C(F)(F)F)[C@H](c2ccc(Cl)cc2)NC(=O)N1

VENDOR NAME: MolPort SC Economical

Explanation of chemical novelty: Maximum Tanimoto similarity to known targets is 0.366.

- Solution 5:

ZINC ID: ZINC000001189603

VENDOR ID: MolPort-002-173-303

SMILES string: CC1=C(C(=O)Nc2ccccc2Cl)[C@@H](c2ccc(C)cc2)NC(=O)N1

VENDOR NAME: MolPort SC Economical

Explanation of chemical novelty: Maximum Tanimoto similarity to known targets is 0.371.

**Submission 9662093**

**Authors: Huiyuan Chen^1^**

**Affiliations: Need affiliation.**

Prediction Methods: |

Assumption; (Guilt by association[1])simlar drugs tend to have similar bahavior on target proteins.

In our method, we first find the compounds (mainly in Drugbank dataset) that can

Bind and inhibit the target provided in the Challenge Question. For example, for Challenge Question 1,

we can find the following the compounds (Drugbank ID) bind their corresponding targets, we called them "existing" compounds;

RET; DB08901 DB08896 DB00398

BRAF; DB08912 DB08881 DB08896 DB00398 DB08881

SRC; DB06616 DB01254 DB09079 DB08901

S6k; None

MKNK1; None

TTK; None

ErK8; None

PAK3; None

PDK1; DB07132 DB07033 DB01933 DB00482 DB04522 DB03777 DB02010

For PDK1, these compounds are not FDA approved, but we still extract their coumpund SMILE string for validation.

And we predict the compounds in two step.

step 1; we want to find the compounds have incitation behavior.

Thus, finding the "novel" compounds that are similar to "existing" compounds for target RET, BRAF, SRC, S6k

step 2; remove the compounds in step 1 that are similar to PDK1's compounds, by doing that we avoid the "novel"

compounds which have anti-targets behavior.

Therefore, we extract other approved compounds (around 2000 compounds, but not include "existing" compounds) in Drugbank,

and measure the compounds similarity by use there chemical structure since chemical properties of a drug are evidently related to its ultimate therapeutic effect

In this challenge, chemical structure of drug in Canonical SMILES form(Simplied Molecular Input Line Entry Specication) are downloaded

from DrugBank. The Chemical Development Kit is then applied to computer the similarity of any two drugs as the Tanimoto

score via their corresponding 2D chemical fingerprints [1]. After measuring the similarity score, we chose the top-5 compounds which have the largest similarity score.

The same processes are applied to Challenge Question 2.

Reference;

1) Altshuler, David, Mark Daly, and Leonid Kruglyak. "Guilt by association." Nature genetics 26.2 (2000); 135.

2) Steinbeck, Christoph, et al. "The Chemistry Development Kit (CDK); An open-source Java library for chemo-and bioinformatics." Journal of chemical information and computer sciences 43.2 (2003); 493-500.

Rationale:

Why is your approach innovative?: Guilt by association is widely used in drug-target prediction, it's simple and effectiveness.

Why will your approach be generalizable?: The approach is generalizable since we only need the "existing" compounds' SMILE strings and measure chemical structure similarity.

Problem 1:

- Solution 1:

ZINC ID: ZINC18516586

VENDOR ID: HY-10571A

SMILES string: CC(C)Nc1cccnc1N1CCN(C(=O)c2cc3cc(NS(C)(=O)=O)ccc3[nH]2)CC1

VENDOR NAME: MedChem Express Economical

Explanation of chemical novelty: we extract the compounds in Drugbank that inhibitor the targets in sub1, then find new compound that are similar to those compounds by their chemical structure.

- Solution 2:

ZINC ID: ZINC150338755

VENDOR ID: HY-15531

SMILES string: CC1(C)CCC(CN2CCN(c3ccc(C(=O)NS(=O)(=O)c4ccc(NCC5CCOCC5)c([N+](=O)[O-])c4)c(Oc4cnc5[nH]ccc5c4)c3)CC2)=C(c2ccc(Cl)cc2)C1

VENDOR NAME: MedChem Express Economical

Explanation of chemical novelty: we extract the compounds in Drugbank that inhibitor the targets in sub1, then find new compound that are similar to those compounds by their chemical structure.

- Solution 3:

ZINC ID: ZINC66166864

VENDOR ID: HY-13011

SMILES string: CCc1cc2c(cc1N1CCC(N3CCOCC3)CC1)C(C)(C)c1[nH]c3cc(C#N)ccc3c1C2=O

VENDOR NAME: MedChem Express Economical

Explanation of chemical novelty: we extract the compounds in Drugbank that inhibitor the targets in sub1, then find new compound that are similar to those compounds by their chemical structure.

- Solution 4:

ZINC ID: ZINC509

VENDOR ID: KS-1086

SMILES string: CN1CCN2c3ncccc3Cc3ccccc3[C@@H]2C1

VENDOR NAME: KeyOrganics Bioactives

Explanation of chemical novelty: we extract the compounds in Drugbank that inhibitor the targets in sub1, then find new compound that are similar to those compounds by their chemical structure.

- Solution 5:

ZINC ID: ZINC896717

VENDOR ID: HY-17492

SMILES string: COc1cc(/C(O)=N/S(=O)(=O)c2ccccc2C)ccc1Cc1cn(C)c2ccc(NC(=O)OC3CCCC3)cc12

VENDOR NAME: MedChem Express Economical

Explanation of chemical novelty: we extract the compounds in Drugbank that inhibitor the targets in sub1, then find new compound that are similar to those compounds by their chemical structure.

Problem 2:

- Solution 1:

ZINC ID: ZINC538658

VENDOR ID: KS-1315

SMILES string: Cc1ccccc1C(=O)Nc1ccc(C(=O)N2CCC[C@@H](O)c3cc(Cl)ccc32)c(C)c1

VENDOR NAME: KeyOrganics Bioactives

Explanation of chemical novelty: we extract the compounds in Drugbank that inhibitor the targets in sub1, then find new compound that are similar to those compounds by their chemical structure.

- Solution 2:

ZINC ID: ZINC20148995

VENDOR ID: HY-B1305

SMILES string: CN(C)CCN(Cc1ccc(Cl)cc1)c1ccccn1

VENDOR NAME: MedChem Express Economical

Explanation of chemical novelty: we extract the compounds in Drugbank that inhibitor the targets in sub1, then find new compound that are similar to those compounds by their chemical structure.

- Solution 3:

ZINC ID: ZINC19632927

VENDOR ID: HY-17037

SMILES string: CN1CCN(CC(=O)N2c3ccccc3C(=O)Nc3cccnc32)CC1

VENDOR NAME: MedChem Express Economical

Explanation of chemical novelty: we extract the compounds in Drugbank that inhibitor the targets in sub1, then find new compound that are similar to those compounds by their chemical structure.

- Solution 4:

ZINC ID: ZINC19632618

VENDOR ID: HY-15463

SMILES string: Cc1ccc(NC(=O)c2ccc(CN3CCN(C)CC3)cc2)cc1Nc1nccc(-c2cccnc2)n1

VENDOR NAME: MedChem Express Economical

Explanation of chemical novelty: we extract the compounds in Drugbank that inhibitor the targets in sub1, then find new compound that are similar to those compounds by their chemical structure.

- Solution 5:

ZINC ID: ZINC6745272

VENDOR ID: HY-10331

SMILES string: CNC(=O)c1cc(Oc2ccc(NC(=O)Nc3ccc(Cl)c(C(F)(F)F)c3)c(F)c2)ccn1

VENDOR NAME: MedChem Express Economical

Explanation of chemical novelty: we extract the compounds in Drugbank that inhibitor the targets in sub1, then find new compound that are similar to those compounds by their chemical structure.
